# AlphaFold Database expands to proteome-scale quaternary structures

**DOI:** 10.64898/2026.03.27.714458

**Authors:** Yewon Han, Maxim I. Tsenkov, Niccolò A. E. Venanzi, Sooyoung Cha, Nilkanth Patel, Sreenath Nair, Razan Abbara, Damian Bertoni, Alejandro Chacón, Nick Dietrich, Boris Fomitchev, Yonathan Goldtzvik, Darren Hsu, Jeannie Austin, Joseph Ellaway, Kieran Didi, Gyuri Kim, Hyunbin Kim, Oleg Kovalevskiy, Dariusz Lasecki, Agata Laydon, Micha Livne, Paulyna Magaña, Maciej Majewski, Urmila Paramval, Risha Patel, Ivanna Pidruchna, Brianda Santini Lopez, Prashant Sohani, Ahsan Tanweer, Duc Tran, Kyle Tretina, Melanie Vollmar, Quan Vu, Augustin Žídek, Sameer Velankar, Martin Steinegger, Jennifer Fleming, Milot Mirdita, Christian Dallago

## Abstract

Protein function is governed by molecular interactions, yet structural coverage of these interactions remains sparse. The AlphaFold Protein Structure Database (AFDB) transformed access to accurate monomeric protein structures at scale. Here, we expand the AFDB to quaternary structures by predicting 31M candidate homo- and heterodimeric protein complexes, compiled from 4,777 proteomes, including model- and global health organisms. We established confidence criteria via analysis of experimentally determined structures, resulting in 1.81M high-confidence predictions. These models enabled the discovery of emergent structures and topologies not present in monomeric predictions. Additionally, the top 1% of structural clusters accounted for ∼44% of all complexes, and ∼8.3% of clusters were conserved across multiple domains of life, pointing to a substantial fraction of ancient, universally retained assemblies. Structural search highlighted that *high-confidence* predictions are anchored in experimental multimer space, yet 31.3% extend beyond detectable PDB coverage. These freely accessible proteome-scale predictions facilitate functional and mechanistic hypothesis generation across biology.

## Introduction

Cellular processes are orchestrated by interacting proteins whose structures encode specificity, regulation, and function^7^. Although interaction networks have been mapped extensively through genetic, biochemical, and computational approaches^2^, the lack of structural information for most protein-protein interactions remains a critical bottleneck to advancing biological understanding, since experimentally determined complex structures in the Protein Data Bank^8^ (PDB) cover only a small fraction of known protein-protein interactions.

The advent of AlphaFold2^9^ enabled rapid and accurate prediction of protein tertiary (3D) structure. The AlphaFold Protein Structure Database (AFDB)^10–12^ provides access to monomeric protein 3D structures across many proteomes. AFDB transformed structural biology and enabled novel basic science discoveries, for instance, the characterisation of novel protein folds^13^, as well as advancing artificial intelligence (AI) in biology, by improving protein 3D prediction methods^14^, or for the development of generative artificial intelligence methods^15^. However, biological function rarely resides in isolated monomers, and for many proteins, biologically relevant conformations, binding sites, and regulatory features only emerge upon complex formation^16,17^. Protein-protein interaction data are available from a range of experimental datasets and are analysed using increasingly sophisticated computational methods. These datasets from diverse biological systems are curated and disseminated through multiple public resources, including the Search Tool for the Retrieval of Interacting Genes/Proteins database (STRING^2^) and IntAct^18^. Translating this interaction knowledge into structural models has the potential to increase the information content available to support mechanistic and functional research.

While comprehensive experimental structural characterisation of all protein interactions remains infeasible, methods predicting structures of protein complexes, such as RoseTTAFold^19^ and AlphaFold-Multimer^4^, have demonstrated that high-confidence quaternary structure prediction is possible. Complex prediction has not yet reached experimental accuracy and still exhibits biases^20,21^, but predicted models can generate strong hypotheses.

Several recent studies characterised proteome-scale protein complex structure predictions in order to generate novel insights into systems biology^22–28^. Yet, these efforts are typically limited to specific organisms, lack consistent confidence calibration, or are not integrated into a stable public infrastructure. As a result, multimer predictions remain difficult to discover, compare, or reuse, preventing their adoption as a component of biological analysis, or in AI training and inference.

Here, we introduce a comprehensive study of protein complex predictions belonging to 4,777 proteomes, including 16 model organisms and 30 proteomes prioritised by the World Health Organization (WHO global health proteomes; **Fig. 1a**), and Swiss-Prot^29^. We present analyses for predictions available as of the 16th of March 2026, comprising 19,148,379 homo- and 7,561,477 heterodimers. We further make 21,456,712 homo- and 7,561,477 heterodimeric predictions available for bulk download, noting that roughly 1.9M candidates failed computational prediction (see “Structure prediction” in Methods; **Supplementary Table 1**). To enable this scale of computation, we extended the acceleration strategies previously applied to monomeric 3D structure prediction^30^ to complex structure prediction, building on AlphaFold-Multimer. Of the analysis-ready set, we identified 1,735,475 homo- and 79,392 heterodimeric high-confidence predictions and integrated them into the AlphaFold Database^4,10^ enabling structural interrogation of interaction networks, variant effects at interfaces, and mechanistic hypotheses that were previously inaccessible at scale. Our work provides a path to improved structure prediction throughput, offers insights into complexes at scale, and provides easy access to novel data through AFDB for systems biology.

**Figure 1:**
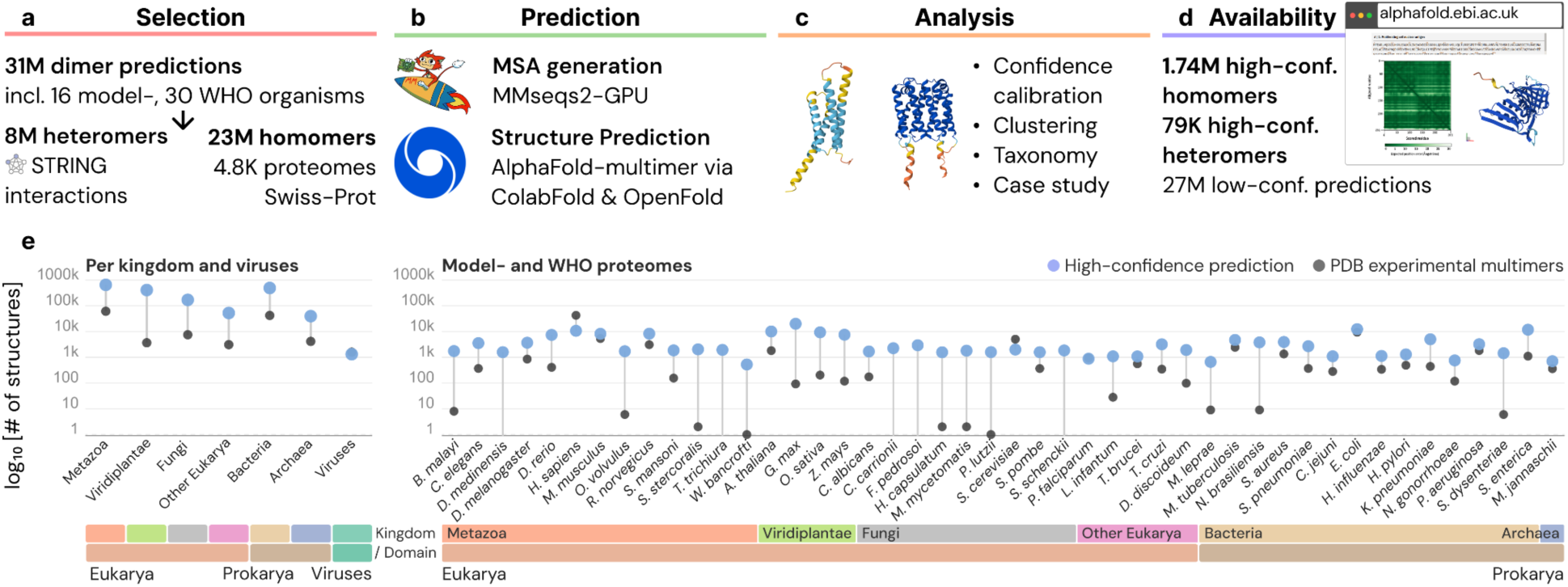
*High-confidence* protein complex 3D predictions extend the AlphaFold Database. **a**, ∼31 million protein homodimer and heterodimer complexes are retrieved from UniProt^1^ and STRING^1,2^. **b**, MSAs are constructed using MMseqs2-GPU^3^, and 3D structures are predicted with AlphaFold-Multimer^4^ through ColabFold^5^ or OpenFold^6^. **c**, We determine reliability threshold and present biologically relevant case studies to highlight emerging behaviour in complex prediction absent from monomer predictions. **d**, High-confidence complex 3D predicted structures are added to the AlphaFold Database for interactive visualisation and interrogation. **e**, Number of high-confidence complexes (blue) and experimentally determined multimers in the PDB (grey), per kingdom and viruses (left) and per model/WHO organism (right).

## Results

We systematically predicted the 3D homodimeric structures for ∼23M annotated sequences derived from 4,777 proteomes in UniProt^1,4,10^ and Swiss-Prot, including 16 model organisms and 30 WHO global health proteomes. Additionally, ∼8M heterodimer candidate pairs were extracted from the “physical protein-protein interaction” set of the STRING Database^2^ by filtering for proteins belonging to the 16 model organisms and 30 WHO global health proteomes (**Fig. 1a**). In contrast to recent studies^22–24^, we decided not to add additional filtering, such as STRING score thresholds, to increase coverage for these critical proteomes. We used MMseqs2-GPU^3^ to generate Multiple Sequence Alignments (MSAs) for homodimers against UniRef100^31^ without the use of metagenomic databases. We established a pragmatic orthology filter that prevents paralogous sequences from diluting the evolutionary signals essential for predicting protein complexes by restricting sequence alignments to the best hit per taxon. This returns monomer alignments, which can be used for homodimer prediction by simply concatenating the alignment with itself row-wise. For heterodimers, we concatenated previously generated MSAs for two different chains without pairing. We then used AlphaFold-Multimer for complex prediction with inference through either an accelerated implementation of ColabFold^5^ or OpenFold^6^ to obtain 3D structures (**Fig. 1b**; Methods).

Once structures were predicted, in addition to model-supplied predicted Local Distance Difference Test (pLDDT) and interface predicted Template Modeling (ipTM) scores, we computed interaction Prediction Score from Aligned Errors (ipSAE)^32^, pDockQ2^33^, and Local Interaction Score (LIS), as well as the number of backbone (clash_backbone_) and heavy atom (clash_heavy-atom_) clashes (**Fig. 1c**). Metrics like ipSAE and pDockQ2 are directional. Therefore we computed scores for chain A relative to chain B and for chain B relative to chain A, and derived *min* and *max* (e.g., ipSAE_min_) values. This reduced directional metrics to a single value per complex independent of direction or cardinality.

Using a benchmark sets of experimentally annotated interacting PDB assemblies and non-interacting entity-pair controls, we identified the combination of ipSAE_max_ greater than or equal to 0.6 and pDockQ2_max_ greater than or equal to 0.23 as high-confidence criteria, resulting in 1,735,475 homodimeric and 79,392 heterodimeric complexes available through the AlphaFold Database with standardised structural formats, confidence metrics, and metadata (**Fig. 1d**).

Overall, this resource provides a massive expansion of the known structural interactome (**Fig. 1e**). The high-confidence predictions consistently exceed the number of experimental multimer structures in the PDB by one to three orders of magnitude across nearly every proteome, with the exception of well-studied organisms *Homo sapiens*, *Escherichia coli* and *Saccharomyces cerevisiae*. Across broader taxonomic clades, particularly *Metazoa* and *Viridiplantae*, these high-confidence models bridge a vast structural gap, providing hypotheses for structural biology at scale.

### Benchmarking identifies high-confidence predictions

To filter predictions for inclusion in AFDB, we benchmarked four protein complex quality metrics (ipTM, ipSAE_max_, LIS_max_, pDockQ2_max_), alongside backbone clash scores to select thresholds that reliably separate PDB annotated physical complexes from non-interacting pairs (**Fig. 2**). We used structurally non-redundant sets of experimental homodimers (n=230) and heterodimers (n=94) released after the AlphaFold-Multimer training cutoff with no detectable dimer-level structural similarity to complexes in the training set, paired with 117 monomeric and 250 non-interacting chain-pair controls. ipSAE_max_ showed the clearest distributional separation and the most stable Matthews Correlation Coefficient (MCC) plateau (**Supplementary Figs. 1,2**), as well as the highest average precision (**Supplementary Fig. 3**), across both homodimer and heterodimer benchmarks. Combining it with pDockQ2_max_ further reduced clash-prone predictions not captured by ipSAE_max_ alone (**Supplementary Fig. 4**). We adopted a combined high-confidence criterion requiring both community-established cutoffs of ipSAE_max_ ≥ 0.6^32,34,35^ and pDockQ2_max_ ≥ 0.23, corresponding to the DockQ ‘acceptable’ quality boundary ^36^. This joint cutoff yielded a precision of 0.924 (False Positive Rate (FPR)= 0.043) for homodimers and 0.958 (FPR=0.004) for heterodimers (**Fig. 2a,b**), supporting its use as a quality filter that prioritises precision over recall given the scale of the release. Applied to the predicted complexes, the filter returned 1,735,475 homodimer (9.1%) and 79,392 heterodimer (1.0%) high-confidence predictions of ∼19.1M predictions selected for analysis (**Fig. 2c,d**). High-confidence predictions also display improved global structural prediction confidence metrics (**Fig 2e,f**; median pLDDT: 62.5 to 87.8 and 66.3 to 80.9; median ipTM: 0.19 to 0.83 and 0.19 to 0.81, respectively). We used max-over min-aggregation across chain directions because ipSAE_min_ is systematically deflated for size-asymmetric heterodimer interfaces, yielding significantly worse benchmark performance (p<0.05; two-sided paired bootstrap test with 2,000 resamples; **Supplementary Fig. 5**); pDockQ2 showed no significant difference, but max-aggregation was retained for consistency, with the joint filter mitigating any added false-positive risk.

**Figure 2.**
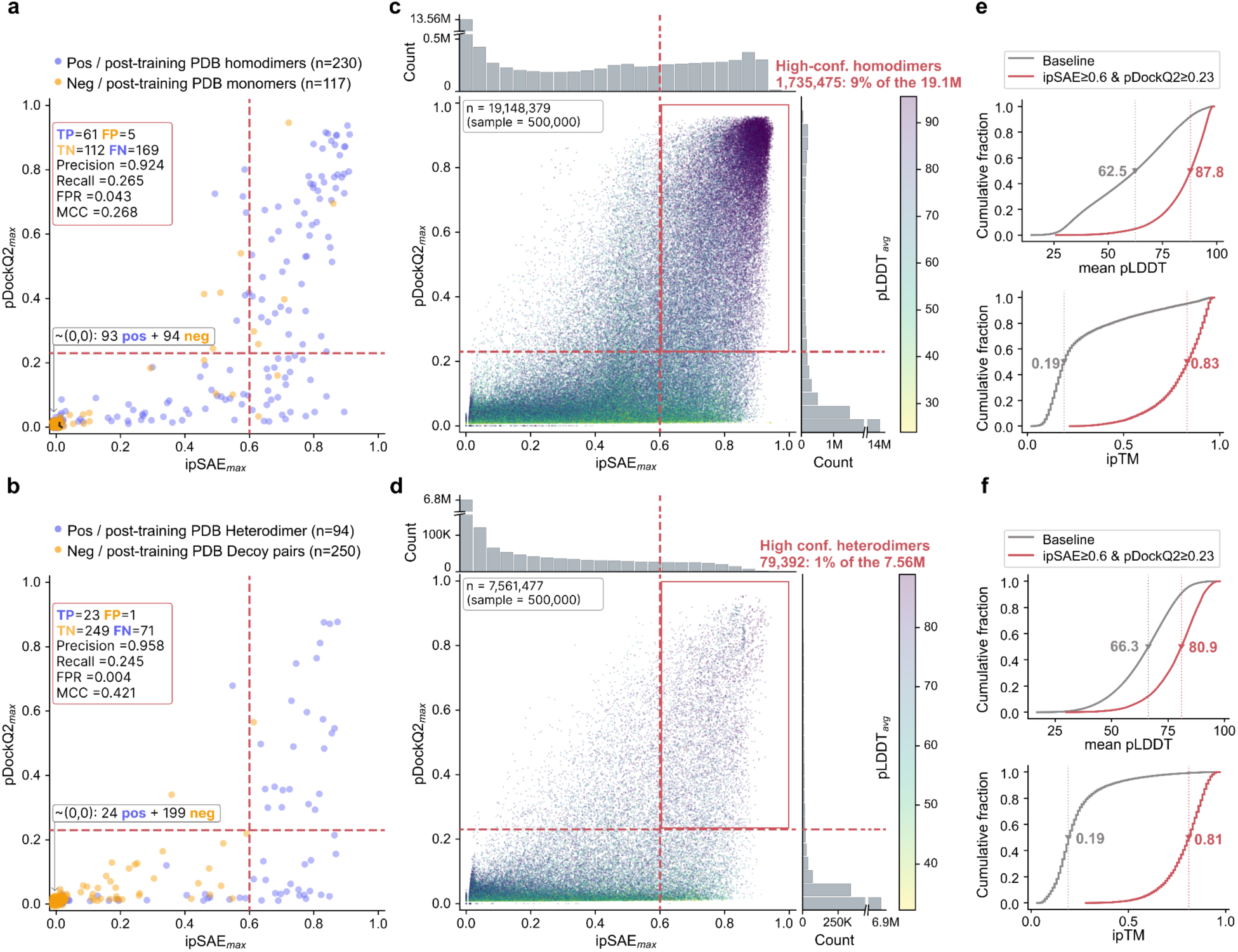
Benchmarking supports joint ipSAE and pDockQ2 thresholds as confidence cutoffs for the selection of high-confidence predicted protein complexes. **a,** Performance of the joint cutoff (ipSAE_max_ ≥ 0.6, pDockQ2_max_ ≥ 0.23) on a homodimer benchmark of structures released after the AlphaFold-Multimer training cutoff (230 positives, 117 monomeric negatives). **b,** As in a, for a heterodimer benchmark (94 positives, 250 not physically interacting chain-pair controls). **c,** Density scatter of ipSAE_max_ versus pDockQ2_max_ for all 19.1 million predicted homodimers, coloured by mean pLDDT. Scatter plots show a random subsample of 500,000 structures; marginal histograms and summary statistics are computed from the full dataset. **d,** As in **c,** for all 7.6 million predicted heterodimers. **e,** CDFs of mean pLDDT (top) and ipTM (bottom) for the full homodimer dataset (grey) and the high-confidence subset (red; n = 1,735,475, 9.1%); triangles mark medians. **f,** As in **e,** for heterodimers (high-confidence subset: n = 79,392, 1.0%). For both homodimers and heterodimers, predictions passing the joint cutoff show substantially higher pLDDT and ipTM.

To test whether similar proteins yield similar dimer predictions, we clustered the high-confidence homodimers at 98% identity using MMseqs2^37^ and aligned each prediction to its cluster representative with Foldseek-Multimer^38^: 94.55% achieved a complex template-modeling score (TM-score) ≥ 0.7, indicating that closely related high-confidence complexes are predicted into similar dimer architectures (**Supplementary Fig. 6a**). Structural alignment of the two chains within each predicted homodimer using Foldseek^39^ further showed that 97.89% of predictions had a TM-score ≥ 0.7 between chains, confirming intra-prediction coherence (**Supplementary Fig. 6b**).

For user interpretation, AFDB entries are additionally labelled by ipSAE_max_ as very high-confidence (≥ 0.8, 982,188 homo- and 23,228 heterodimers), confident (0.7–0.8, 440,072 homo- and 31,728 heterodimers), or low-confidence (0.6–0.7, 313,215 homo- and 24,436 heterodimers).

### Determinants of high-confidence complex predictions

By taxonomy, Archaea and Bacteria exhibit the highest high-confidence fractions for both homo-(∼27% of all homodimer predictions) and heterodimers (∼2–3% of all heterodimer predictions), while most eukaryotic kingdoms fall below the overall baseline (9.1%), with the exception of Fungi, which exceeds it for homodimers (11.2%; **Fig. 3a, b**). This pattern is consistent with the lower prevalence of long disordered regions in prokaryotic proteomes, which favours recovery of stable oligomeric interfaces, in line with the successful proteome-wide prediction of homo-oligomers in an archaeon reported previously^26^. High-confidence homodimers are enriched at intermediate chain lengths, peaking at ∼330–410 residues, with lower confidence at both shorter and longer extremes (**Fig. 3c**). This trend holds for heterodimers as well (**Supplementary Fig. 7a**). Additionally, heterodimeric pairs with similar chain lengths also showed a higher high-confidence fraction, suggesting high-confidence correlation with homodimer-likeness. However, when stratified by sequence and length similarity (**Supplementary Fig. 7b**), these homodimer-like pairs represented only a small subset of the full heterodimer set, thus most predicted STRING-derived heterodimers are unlikely to be driven by homodimer-like pairing.

**Figure 3:**
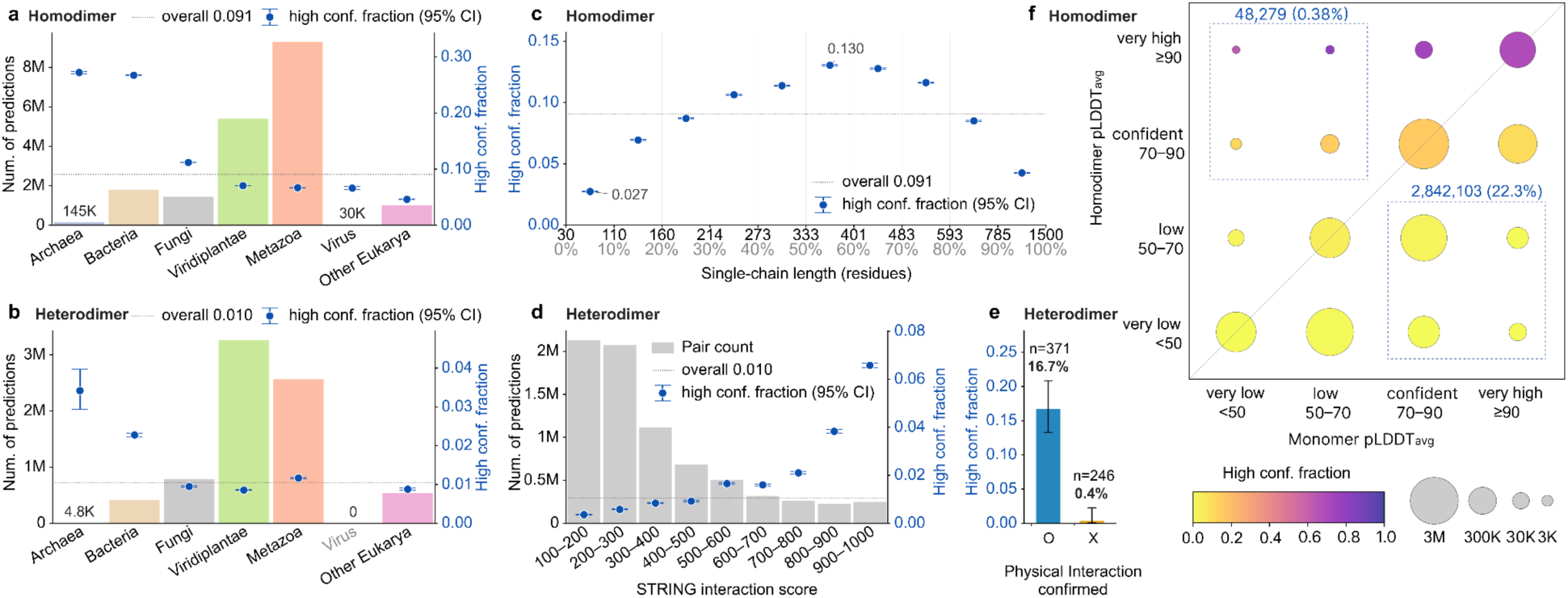
Factors determining high-confidence homo- and heterodimeric predictions. **a, b,** Per-kingdom high-confidence fraction for homodimers (a; 1.74 M high-confidence of 19.1 M total) and heterodimers (b; 79.4 K high-confidence of 7.56 M total), defined as ipSAE_max_ ≥ 0.60 and pDockQ2_max_ ≥ 0.23. Coloured bars (left y-axis), per-kingdom input pool size; navy dots (right y-axis), per-kingdom high-confidence fraction; dashed lines, overall high-confidence fractions (0.091 and 0.010, respectively); whiskers, 95% Wilson confidence intervals. **c,** Homodimer high-confidence fraction across single-chain length deciles. Top x-axis, residue cutoffs; bottom, cumulative percentiles. **d,** Heterodimer high-confidence fraction across STRING physical-interaction score bins (width 100). Grey bars, pair count (left axis); blue dots, high-confidence fraction (right axis). **e,** Fraction of high-confidence predictions among post-training STRING-derived heterodimer pairs with the direct chain contact confirmed by the experimental structures from PDB (n = 371) or no direct contact (n = 246). **f,** Joint distribution of AFDB v6 monomer and homodimer pLDDT_avg_ for 12.8 M sequence-matched predictions on a 4 × 4 confidence-category grid. Bubble area is proportional to square root of the count; colour encodes within-cell high-confidence fraction. Blue dotted boxes highlight two contrasting regimes: upper-left, predictions with confident-to-very-high dimer pLDDT but low-to-very-low monomer pLDDT; lower-right, the converse.

For heterodimers, the high-confidence fraction scales with STRING physical-interaction evidence, rising approximately 18-fold from the lowest to the highest score bin (**Fig. 3d**). STRING-derived heterodimer pairs with direct chain-chain contact in post-training PDB structures showed a higher fraction of high-confidence predictions than the non-contacting chain-pair controls (**Fig. 3e**).

Finally, we compared monomer and dimer pLDDT for ∼12.8 M homodimers with exact-sequence matches in AFDB v6 (**Fig. 3f**). Predictions in which dimer pLDDT markedly exceeds monomer pLDDT (upper-left box) are enriched for high-confidence interface scores relative to the baseline (9.8% for the 12.8 M sequence-matched predictions), suggesting that oligomerisation can stabilise regions that are poorly predicted as monomers for this small fraction. A similar enrichment in the upper-left box is observed among AFDB v6-matched heterodimer pairs (**Supplementary Fig. 8**). Furthermore, this indicates that, for a subset of proteins, oligomeric context provides structural constraints absent from monomer prediction, allowing recovery of folds or interfaces that remain weakly specified in monomeric models (**Fig. 4**).

**Figure 4:**
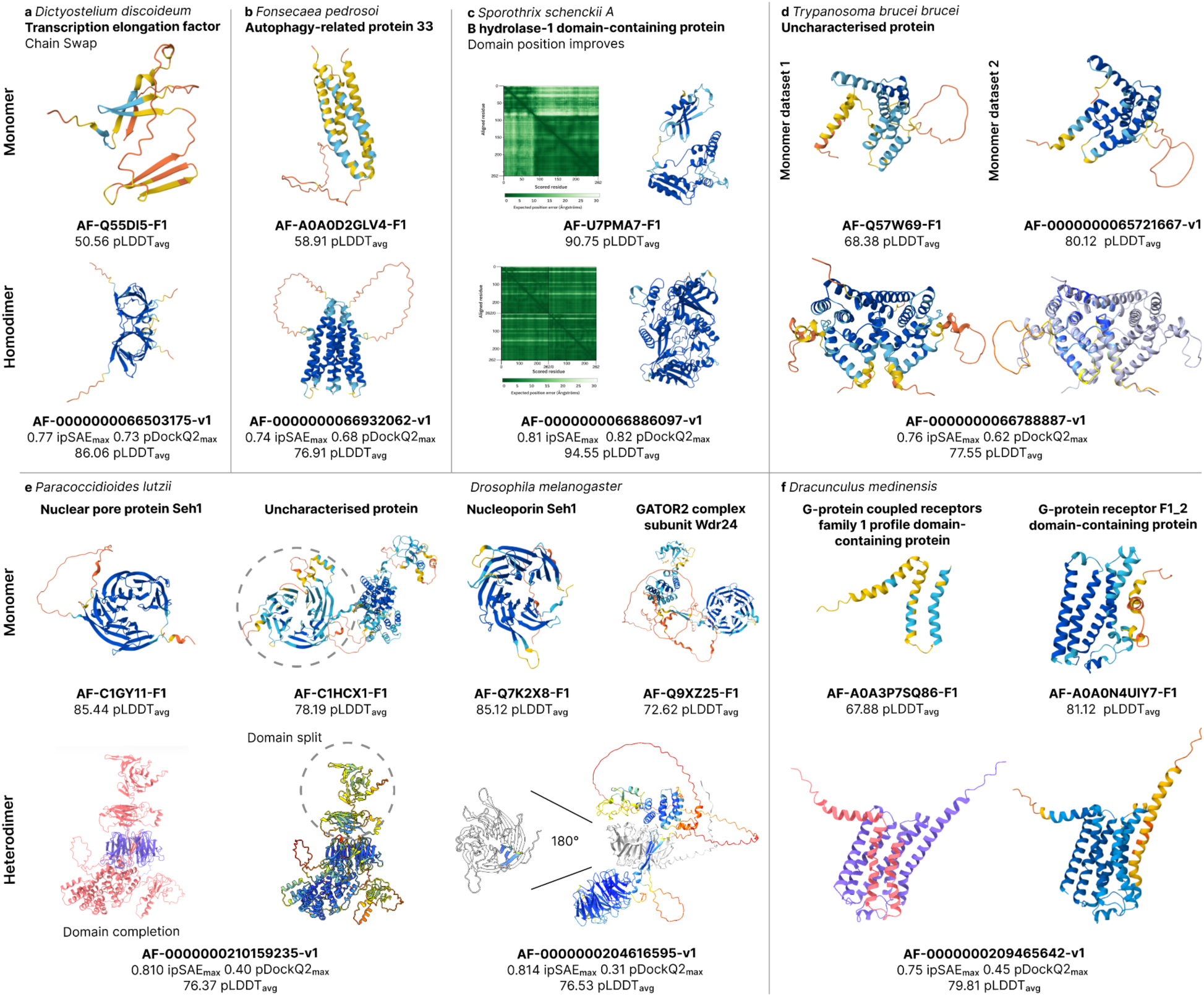
Oligomeric context influences the structural interpretation of proteins. **a**, For proteins like Transcription elongation factor Eaf N-terminal domain-containing protein (UniProt accession Q55DI5), a high-confidence fold emerges only in the homodimer prediction through domain swapping, a structure entirely missed by the low-confidence monomeric model. **b**, Similarly, for the membrane autophagy-related protein 33 (UniProt accession A0A0D2GLV4), the dimeric model yields a more coherent, high-confidence assembly that better defines membrane boundaries. **c**, Multimer prediction can refine the global inter-domain architecture even for an already confident monomeric model, such as an AB hydrolase-1 domain-containing protein (UniProt accession U7PMA7). **d**, For *Trypanosoma brucei brucei* UniProt accession Q57W69, both MSA quality and oligomeric context can improve the resulting model confidence in the same manner. **e**, Heteromeric prediction can also recover composite domain architectures. In *Paracoccidioides lutzii*, an uncharacterised protein (UniProt accession C1HCX1) contributes three β-strands that complete the WD40 β-propeller of the nuclear pore protein Seh1 (UniProt accession C1GY11), a feature absent from the monomeric Seh1 model despite high overall confidence. A similar Seh1-complementing interaction is observed in *Drosophila melanogaster* between Seh1 (UniProt accession Q7K2X8) and the GATOR2 subunit Wdr24 (UniProt accession Q9XZ25). **f**, Hetero-oligomeric modelling can stabilise helical assemblies that remain incomplete in monomeric predictions. In *Dracunculus medinensis*, G-protein coupled receptors family 1 profile domain-containing protein (UniProt accession A0A3P7SQ86) forms only a small three-helix bundle in isolation, but assembles with G_PROTEIN_RECEP_F1_2 domain-containing protein (UniProt accession A0A0N4UIY7) into a substantially more ordered six-helix architecture with increased overall confidence.

### Oligomeric context reshapes protein structure

To illustrate the range of ways in which oligomeric context can affect structural interpretation, we selected representative examples drawn from eukaryotic microbes, fungal and bacterial pathogens, a neglected tropical parasite, and a crop species. These examples were chosen to capture different outcomes, including recovery of domain-swapped folds, improved membrane-protein organisation, refinement of inter-domain architecture, and agreement with models derived from curated sequence input (**Fig. 4**).

The transcription elongation factor Eaf N-terminal domain-containing protein from *Dictyostelium discoideum* (UniProt accession Q55DI5) provides a clear example of a fold that emerges only in an oligomeric context. The monomeric model (AF-Q55DI5-F1) has low confidence (pLDDT_avg_=50.56) and fragmented β-sheet elements. In contrast, modelling the protein as a homodimer (AF-0000000066503175-v1) yields a well-defined structure (pLDDT_avg_=86.06) formed by domain swapping between the two chains (**Fig. 4a**). Each chain contributes structural elements that complete the fold of its partner, such that the architecture is assembled across chains rather than within a single polypeptide. A Foldseek search against the PDB identified a related architecture in the structure of an active transcription elongation complex Pol II-DSIF (SPT5-KOW5)-ELL2-EAF1 (pdb_00007okx), in which the fold is likewise formed across chains O and M, supporting the domain-swapped assembly observed in the prediction.

The Autophagy-related protein 33 from *Fonsecaea pedrosoi* (UniProt accession A0A0D2GLV4) provides a membrane-protein example in which oligomeric context improves structural definition. The monomeric model already contains a four-helix bundle, but with relatively low confidence (pLDDT_avg_=58.91). In contrast, the dimeric model brings two such bundles together into a much more coherent high-confidence assembly (pLDDT_avg_=76.91, ipSAE_max_=0.74), and more clearly defines the likely membrane boundaries, with the lower-confidence regions (light blue) lying mainly outside the membrane-spanning layer (**Fig. 4b**). Measuring the high-confidence stretches of the transmembrane helices gives distances of 34-39 Å, which is similar to the thickness achieved by the lipids packing inside a lipid bilayer (∼30-40 Å), i.e. the main component of cell membranes. This case suggests that, for some membrane proteins, monomeric prediction can recover the core topology, whereas oligomeric modelling is needed to better resolve the full assembly.

The monomeric AlphaFold2 model of *Sporothrix schenckii* AB hydrolase-1 domain-containing protein (AF-U7PMA7-F1) is already highly confident overall (**Fig. 4c**), with pLDDT_avg_=90.75, indicating that the local fold is well specified. However, modelling the protein as a dimer improved the relative positioning of the domains, as seen in the reduced uncertainty in the Predicted Aligned Error (PAE) plot, suggesting that oligomeric context helps constrain the inter-domain arrangement rather than the local secondary structure itself. This illustrates that even when a monomeric model is confident, multimeric prediction can still refine the global architecture and identify interacting interfaces.

For the uncharacterised *Trypanosoma brucei brucei* protein (UniProt accession Q57W69), different modelling inputs produced materially different structural hypotheses (**Fig. 4d**). The original model showed only moderate confidence (pLDDT_avg_=68.38), whereas use of a curated multiple sequence alignment, following the trypanosomatid-focused approach of Wheeler et al.^40^, improved the prediction to pLDDT_avg_=80.12. The dimeric model agrees closely with this improved dataset-derived structure: superposition of AF-0000000066788887-v1 chain A with AF-0000000065721667-v1 chain A gives a Root Mean Square Deviation (RMSD) of 0.67 Å over 166 pruned atom pairs. This agreement between the curated-MSA and dimeric models indicates that both MSA composition and oligomeric state influence the resulting prediction.

Similar patterns of improvement can be seen within the heteromeric complexes. In the fungal pathogen *Paracoccidioides lutzii* (*P. lutzii)*, a predicted complex between the nuclear pore protein Seh1 (UniProt accession C1GY11) and an uncharacterised protein (UniProt accession C1HCX1) resolves a gap in the Seh1 WD40 β-propeller that remains incomplete in the monomeric model despite high overall confidence (pLDDT_avg_=85.44) (**Fig. 4e**), with three β-strands from the uncharacterised partner C1HCX1 completing the missing blade architecture. This arrangement resembles experimentally characterised Seh1-containing complexes in which partner proteins create composite β-propeller domains (nine examples of experimental Seh1 complexes are available in humans^41^; eleven in yeast^42^). Although C1HCX1 lacks functional annotation, its STRING partners include SEC13-family transport protein (UniProt accession C1H9N1), WD repeat-containing proteins (UniProt accession C1H7P7 and C1GNV8), and nitrogen permease regulator Npr2 (UniProt accession C1GR16). These predicted dimers did not pass the high-confidence threshold, indicating limited support for stable binary interactions and leaving open whether C1HCX1 acts through higher-order assemblies, or whether the predicted partners represent indirect or non-physical associations. The improvement in the complex is local rather than global. The Seh1 interaction interface is well resolved, whereas the WD40-repeat region UniProt accession of C1HCX1, not involved in the Seh1 interaction, separates into two lobes and shows reduced confidence relative to the monomeric prediction (**Fig. 4e**, grey dashed circle), illustrating how local interaction features can remain coherent even when confidence deteriorates elsewhere in a multidomain assembly. Interestingly, a similar assembly principle is observed in *Drosophila melanogaster*, where the WD40-repeat nucleoporin Seh1 (UniProt accession Q7K2X8) forms a predicted interaction with the GATOR2 complex subunit Wdr24 (UniProt accession Q9XZ25), another WD40-repeat-containing protein involved in TOR signalling regulation. As in *P. lutzii*, the WD40 β-propeller region of Seh1 is completed by its complexation partner. The similarity between the fungal and fly complexes (as well as the experimental human and yeast complexes) is striking, revealing a conserved peptide-mediated assembly principle and higher-order scaffold architecture across the tree of life.

In the Guinea worm parasite *Dracunculus medinensis*, a major target of ongoing global eradication efforts, we predicted a membrane-protein assembly between an uncharacterised G-protein coupled receptors family 1 profile domain-containing protein (UniProt accession A0A3P7SQ86) and another G-protein receptor protein (UniProt accession A0A0N4UIY7; **Fig. 4f**). Individually, A0A3P7SQ86 is predicted with relatively low confidence (pLDDT_avg_=67.9), and forms only a small three-helix bundle, whereas A0A0N4UIY7 is predicted with higher overall confidence (pLDDT_avg_=81.1) except for a low-confidence 40 residue C-terminal region. When modelled together, the two proteins assemble into a coherent six-helix architecture. A0A3P7SQ86 packs closely against the helices of A0A0N4UIY7, increases in confidence for both components (pLDDT_avg_=74.4 for A0A0N4UIY7 and pLDDT_avg_=93.2 for A0A3P7SQ86), producing a consistent, likely membrane-spanning bundle due to their length (∼30 Å) and hydrophobic nature. The assembly as a whole reaches a pLDDT_avg_ of 79.8, with the improvement concentrated in A0A3P7SQ86; the low-confidence C-terminal tail of A0A0N4UIY7 remains as a low-confidence region. As in the preceding examples, oligomeric context does not uniformly improve all regions of the model, but instead stabilises specific structural features that remain incomplete or weakly resolved in isolation. Here, the principal effect is the organisation of transmembrane helices into a shared membrane assembly, consistent with a fold that depends on partner association within the membrane environment.

### Predicted complexes connect to and extend beyond experimental multimers

High-confidence homodimeric and heterodimeric structures were clustered at the complex level using Foldseek Multimercluster (manuscript in preparation), considering both chain and interface structural similarity. This compressed 1,814,832 structures 5.4-fold into 333,787 clusters, of which 90,388 contained at least one other member (non-singleton clusters; **Fig. 5a**). At the individual level, 568,302 high-confidence complexes (31.3%) lacked detectable PDB multimer analogs under our Foldseek multimer search criteria (Methods); at the cluster level, this fraction rises to 63.6% (212,351 of 333,787 representatives; **Fig. 5a**). Conversely, 68.7% of individual complexes and 36.4% of clusters match at least one PDB multimer, indicating that the high-confidence set is structurally anchored in known structural space, while expanding substantially beyond it.

**Figure 5.**
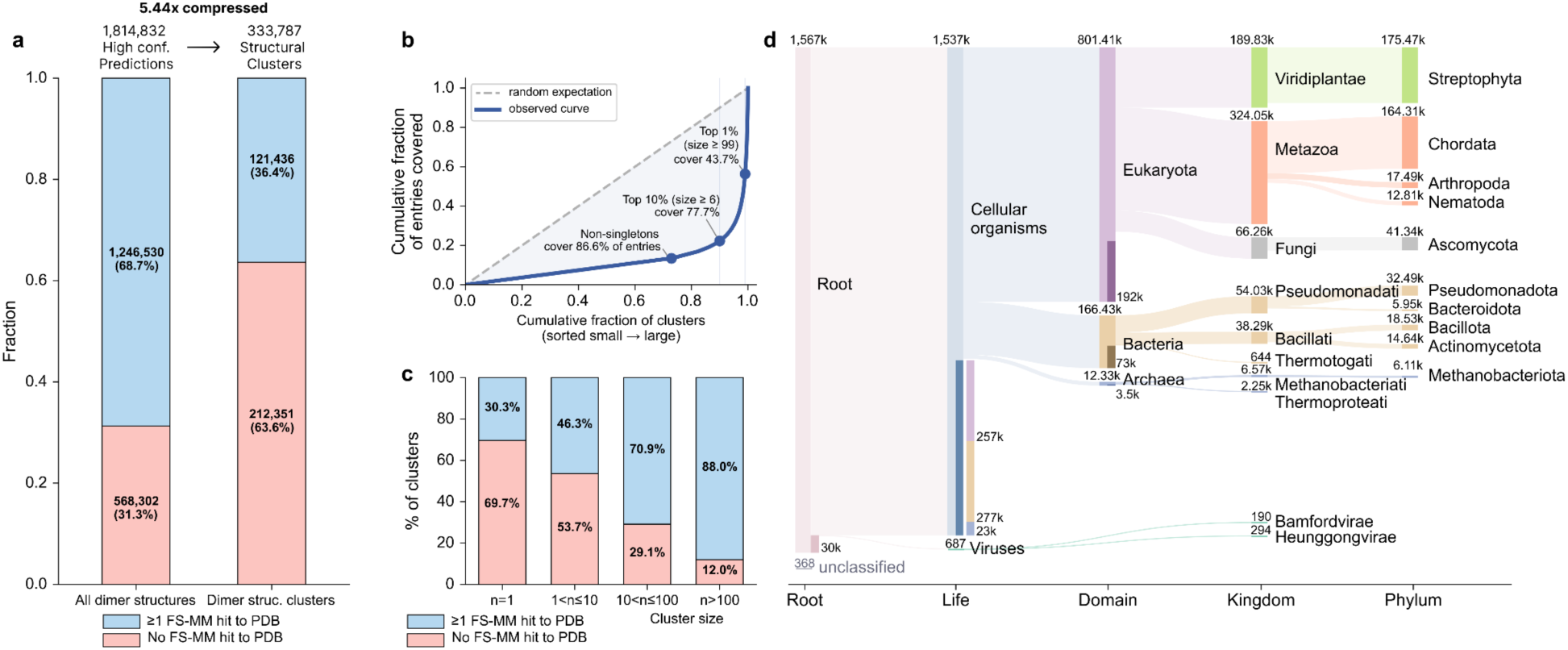
Structural clustering of high-confidence complexes reveals novelty beyond experimental structures and conservation across the tree of life. **a,** Proportion of structures with at least one Foldseek-Multimer (FS-MM) structural search hit to PDB, before and after clustering. Of 1,814,832 high-confidence (ipSAE_max_ ≥ 0.6 and pDockQ2_max_ ≥ 0.23) complexes available for structure-based analysis, 68.7% have a matching PDB structure. After clustering, 36.4% of 333,787 clusters contain at least one member with a PDB hit. **b,** Lorenz curve showing the cumulative fraction of entries covered as a function of the cumulative fraction of clusters, sorted from smallest to largest. The observed curve (solid blue) deviates from the random expectation (dashed diagonal), indicating that a small number of large clusters account for a disproportionate share of entries. Non-singleton clusters (27.1% of all clusters) cover 86.6% of entries, and the top 1% of clusters (size ≥ 99) alone account for 43.7%. **c,** Percentage of clusters containing at least one PDB hit, stratified by cluster size. Larger clusters are more likely to contain a PDB-matched member, with 88.0% of clusters with size > 100 having at least one hit, compared to 30.3% for singletons. **d,** Sankey^43^ diagram of complexes in non-singleton clusters across taxonomic ranks. Each complex is labelled by its cluster’s lowest common ancestor. Bands in the Root, Life, and Domain columns show complexes whose LCA sits exactly at that rank; in the “Cellular organisms” node, these are further split by member domain.

The cluster size distribution was highly skewed: the largest 1% and 10% of clusters encompassed 43.7% and 77.7% of all predicted complexes, respectively (**Fig. 5b**). Singletons account for only 13.4% of entries despite making up 73% of clusters, indicating that predicted high-confidence complex space is concentrated around a small number of recurrent complex architectures. In addition, non-singleton clusters that display no detectable PDB multimer match were more frequent among smaller clusters than among the largest clusters (**Fig. 5c**), suggesting that structurally recurrent solutions are more likely to overlap with known multimeric space, whereas structurally rarer clusters are more often unsupported by current PDB coverage.

Clusters containing only homodimers accounted for 89.8% of non-singleton clusters; heterodimer-only and mixed accounted for 8.14% and 2.04% respectively. Heterodimers in mixed clusters showed markedly higher inter-chain TM-score than those in hetero-only clusters (median, 0.91 vs. 0.12), supporting their interpretation as predominantly homodimer-like **(Supplementary Fig. 9)**.

Across the top 1% of cluster representatives (n = 3,338), 49.1% carry a UniProt Enzyme Commission (EC) annotation, covering 55.2% of their members, with transferases (36.0%) and oxidoreductases (21.6%) the leading EC classes. Eight of the ten largest clusters are annotated with a single EC class (**Supplementary Fig. 10**). The largest cluster (1,404 members; majority EC 1.1.1.284) corresponds to alcohol dehydrogenases. This stands in contrast to a prior homo-oligomer atlas that covered four proteomes and identified ∼3,000 dimer structure types dominated by coiled-coil-mediated dimerisation^26^, illustrating how scale shifts the dominant recurrent structures from coiled-coil motifs to conserved catalytic families.

To assess evolutionary conservation, we computed the lowest common ancestor (**Fig. 5d**) for each cluster. Notably, ∼8.3% of non-singleton clusters contain members from at least two different domains, that is, with the lowest common ancestor (LCA) at the level of cellular organisms (n = 7,194) or above (n = 305), and these collectively cover ∼32.3% of all predicted complexes (586,068 of 1,814,832). This suggests that a substantial fraction of high-confidence complexes may correspond to deeply conserved quaternary molecular architectures shared across cellular life.

## Discussion

This work extends AFDB from monomeric structural coverage to quaternary structures at proteome scale. To realise this, we compiled over 31 million candidate complexes and surfaced 1.81 million high-confidence assemblies. These span reference proteomes, Swiss-Prot, and WHO-prioritised organisms, including neglected-disease pathogens. Integration into AFDB facilitates search and interpretation, turning large-scale complex prediction into a durable community resource.

We found that oligomeric context does not only increase model confidence, but can alter how the structure is interpreted. In representative cases, complex prediction resolves domain-swapped folds, membrane assemblies, inter-domain arrangements, and composite domain architectures that are weakly specified or absent in monomeric models. These results underscore that, for a subset of proteins, the biologically relevant structural unit is the assembly rather than the isolated chain. Complex search and clustering show that predicted complex space connects to- and extends beyond experimental multimers. Recurrent predicted families are frequently anchored in known multimeric space, whereas many cluster representatives lack detectable PDB analogs under our criteria. As homodimer and heterodimer prediction expands across additional proteomes, especially underrepresented taxa, some singleton clusters may resolve into broader recurrent families.

Our joint ipSAE and pDockQ2 cutoff for high-confidence predictions is supported by a benchmark designed to reduce training-set overlap at the quaternary assembly structure level rather than at the single-chain level. This joint cutoff for high-confidence predictions prioritises confidence for AFDB integration; below-threshold predictions remain available for download and may still contain true interactions.

The high-confidence fraction is lower for heterodimers (1.0%) than for homodimers (9.1%), likely attributable to two reasons: 1) heterodimer candidates were drawn from STRING physical-interaction annotations without score filtering, so many candidate pairs carry weak interaction evidence. 2) Physical association from STRING does not necessarily imply direct/dimer contact, which we observed (**Fig. 3e)**.

Together, these results establish a scalable framework for mapping protein assembly space and prioritising experimentally uncharacterised interfaces across diverse proteomes. We anticipate that this resource will facilitate large-scale analyses of protein-protein interfaces and can serve as a structural foundation for applications in drug discovery, protein engineering, and the training of next-generation biological AI models.

## Availability

1,735,475 high-confidence homodimer predictions and 79,392 high-confidence heterodimers, selected using the criteria ipSAE_max_ ≥ 0.6, pDockQ2_max_ ≥ 0.23, are available as individual entry pages through AlphaFold Database at alphafold.ebi.ac.uk. To support AFDB users in interpreting the structure models, surfaced entries are further categorised in “*very high-confidence”* (ipSAE_max_ ≥ 0.8), “*confident”* (0.7 ≤ ipSAE_max_ < 0.8), and “*low-confidence*” (0.6 ≤ ipSAE_max_ < 0.7). Dimers not passing the previously defined threshold, together with their interface scores, are provided on the FTP page ftp.ebi.ac.uk/pub/databases/alphafold/collaborations/nvda/. These models and MSAs are also available through bulk download to support large-scale computational analysis, benchmarking, and method development, while reducing the risk of over-interpretation in routine biological use.

ColabFold with improved throughput is available at github.com/sokrypton/ColabFold in release 1.6.0. Acceleration libraries cuEquivariance (docs.nvidia.com/cuda/cuequivariance) and TensorRT (docs.nvidia.com/deeplearning/tensorrt), that were used to improve OpenFold throughput, are freely available with Apache 2.0 licensing. OpenFold with TensorRT and cuEquivariance integration is available at github.com/aqlaboratory/openfold.

## Acknowledgements

We would like to thank NVIDIA colleagues, in particular Kyle Gion, Christian Hundt, Tobias Lasser, Isabel Wilkinson, Anthony Costa, Xin Yu, Hari Sadasivan, Youhan Lee, and Yuxing Peng for support. EMBL-EBI also acknowledges their colleagues Oana Stroe, Gemma Wood, and Victoria Hatch for their support. Martin Steinegger acknowledges support by the National Research Foundation of Korea (NRF) grants funded by the Korea government (MSIT) (2020M3A9G7103933, RS-2021-NR061659 and RS-2021-NR056571, RS-2024-00396026), Novo Nordisk Foundation (NNF24SA0092560), and Creative-Pioneering Researchers Program through Seoul National University. Milot Mirdita acknowledges support from the National Research Foundation of Korea (NRF) grants funded by the Korea government (MSIT) (RS-2023-00250470, RS-2026-25515455). This work was supported by the BBSRC, UK Research and Innovation [20-BBSRC/NSF-BIO], Google DeepMind and the European Molecular Biology Laboratory. The AlphaFold Protein Structure Database is supported by Google DeepMind and the European Molecular Biology Laboratory.

## Contributions

Razan Abbara (Validation), Damian Bertoni (Software, Validation), Jennifer Fleming (Conceptualisation, Validation, Formal analysis, Supervision, Project administration, Writing - Original Draft, Writing - Review & Editing), Yonathan Goldtzvik (Validation), Sreenath Nair (Methodology, Software, Investigation, Formal analysis, Data Curation, Supervision), Paulyna Magaña (Validation), Urmila Paramval (Validation, Software), Ivanna Pidruchna (Validation, Software), Ahsan Tanweer (Investigation, Methodology, Data Curation, Validation, Software), Maxim I. Tsenkov (Investigation, Software, Validation, Formal analysis, Data Curation, Methodology), Joseph Ellaway (Validation, Software, Data Curation), Jeannie Austin (Project administration), Melanie Vollmar (Validation, Formal analysis), Niccolò A. E. Venanzi (Software, Validation, Formal analysis, Investigation, Methodology, Data Curation), Darren Hsu (Software, Validation, Data Curation, Methodology), Nilkanth Patel (Software, Validation, Data Curation, Methodology), Alejandro Chacón (Software, Validation), Duc Tran (Software, Validation), Quan Vu (Software), Boris Fomitchev (Software), Brianda Santini Lopez (Software, Validation, Data Curation), Kyle Tretina (Writing - Original Draft, Writing - Review & Editing), Maciej Majewski (Software, Validation, Methodology, Investigation, Formal analysis), Kieran Didi (Formal analysis), Gyuri Kim (Visualisation), Hyunbin Kim (Visualisation, Writing - Review & Editing), Micha Livne (Data Curation), Prashant Sohani (Data Curation), Yewon Han (Methodology, Software, Validation, Investigation, Formal analysis, Data Curation, Visualisation, Writing - Original Draft, Writing - Results & Methods, Writing - Discussion, Writing - Review & Editing), Sooyoung Cha (Validation, Investigation, Formal analysis, Visualisation, Writing - Original Draft, Writing - Methods), Milot Mirdita (Conceptualisation, Methodology, Software, Investigation, Formal analysis, Data Curation, Supervision, Writing - Review & Editing), Christian Dallago (Conceptualisation, Methodology, Software, Validation, Formal analysis, Investigation, Resources, Data Curation, Writing - Original Draft, Writing - Review & Editing, Supervision, Project administration, Funding acquisition),

Martin Steinegger (Conceptualisation, Methodology, Software, Validation, Investigation, Formal analysis, Resources, Data Curation, Supervision, Project administration, Funding acquisition, Writing - Original Draft, Writing - Review & Editing), Sameer Velankar (Conceptualisation, Methodology, Software, Data Curation, Validation, Investigation, Resources, Supervision, Project administration, Funding acquisition, Writing - Review & Editing), Nick Dietrich (Writing - Review & Editing), Oleg Kovalevskiy (Writing - Review & Editing), Dariusz Lasecki (Writing - Review & Editing), Agata Laydon (Writing - Review & Editing), Risha Patel (Writing - Review & Editing), Augustin Žídek (Writing - Review & Editing)

## Methods

### Sequence and interaction dataset construction

#### Sequence selection and interaction definition

We selected 23,441,822 UniProt 2025_04 sequences by filtering UniProt for a set of proteomes and for proteins with length between 15 and 1,500 amino acids. The set of 4,777 proteomes was selected from Swiss-Prot, proteomes within the World Health Organization (WHO) global health proteomes, and the top downloaded reference proteomes in UniProt. From this set, homodimers were simply derived by duplicating each monomer into a complex, thus resulting in 23,441,822 homodimers, with the longest homodimers in the set containing a maximum of 3,000 amino acids.

Heterodimers were instead derived by extracting physical interaction evidence provided by the STRING database. In particular, the file protein.physical.links.v12.0.txt.gz (11.1 GB) CC-BY-4.0, obtained from https://string-db.org/cgi/download in January of 2026, was downloaded. This file contains annotations of physically interacting partners with their corresponding STRING score. From these interactions, we filtered for sequences within a prioritised set of proteomes from 16 model organisms and 30 global health-relevant proteomes, resulting in 7,620,644 candidate complexes with two distinct chains. Further, these interactions were filtered to produce heterodimers of maximally 3,000 amino acids. No further filtering, such as STRING score thresholds, were applied, to obtain the highest coverage.

#### STRING ID to UniProt accession mapping

To map STRING (v12.0) physical interaction proteins to our UniProt proteomes, we applied a three-step strategy with decreasing priority. (1) UniProt ID mapping: STRING provides a direct STRING-to-UniProt identifier mapping via the file protein.aliases.v12.0.txt.gz (CC-BY-4.0), downloaded from https://string-db.org/cgi/download in January 2026. As this file contains obsolete UniProt accessions, we re-mapped all entries against the UniProt 2025_04 release, obtained from https://ftp.ebi.ac.uk/pub/databases/uniprot/current_release/knowledgebase/complete/ in January 2026. (2) CRC64 hash matching: For proteins not resolved by step 1, we matched CRC64 sequence hashes between STRING and UniProt entries. (3) MMseqs2 sequence search: For the remaining unmapped proteins, we ran per-taxon MMseqs2 searches (STRING sequences as queries against our reference set) requiring 100% sequence identity and ≥95% bidirectional coverage (--min-seq-id 1 -c 0.95 --cov-mode 0 --alignment-mode 3 -s 4), retaining only the best hit per query based on E-value and bit score.

For both the ID mapping and CRC64 steps, we applied a taxon priority filter: when a STRING protein could be mapped to multiple UniProt accessions, same-species mappings (where the taxon ID extracted from the STRING identifier matched the UniProt taxon ID via the NCBI taxonomy dump; downloaded in March 2026) were preferred; cross-taxon mappings were retained only when no same-species match existed. When multiple methods successfully mapped the same STRING protein, the highest-priority result was kept (ID mapping > CRC64 > MMseqs2).

Finally, STRING physical interaction pairs were deduplicated such that each unordered pair (A–B / B–A) appeared only once, and the mapped UniProt accessions were converted to internal identifiers (AF-series IDs) using a reference table, producing per-taxon interaction files. This pipeline achieved a mapping rate of 73.1% for the model organism set (5,503,385 out of 7,528,643 STRING entries) and 82.4% for the WHO global health proteome set (2,145,870 out of 2,604,057). Redundancy removal resulted in 7,620,644 candidates for heterodimeric prediction.

### Inference tools and approaches

To improve computational efficiency, we separated the two compute-demanding workloads of MSA generation and structure prediction.

#### MSA generation

We generated homodimer MSAs by leveraging a pre-release version of ColabFold 1.6.0 using the MMseqs2-GPU backend (version 18-8cc5c). In particular, we leveraged the *colabfold_search* tool with a new MSA filtering strategy for homomeric predictions. This strategy keeps only the best hit per taxon based on alignment score, essentially returning a monomer MSA in a3m format filtered to contain only the highest scoring hit per taxon. The MSA is then duplicated during AlphaFold-Multimer’s featurisation for homodimeric prediction. *colabfold_search* was run using the following parameters: --use-env 0 --pairing_strategy 1 --pair-mode paired --filter 2 --db-load-mode 2. The underlying sequence database was the pre-built ColabFold GPU-accelerated search database uniref30_2302, which provides clustered coverage of all UniRef100 sequences.

To generate heterodimer MSAs, we simply concatenated the previously obtained homodimer/monomer MSAs for two distinct sequences, without additional pairing. This design was chosen on the one hand, to decrease computational complexity, and on the other, as these input MSAs were already restricted to the best hit per taxon, the need for pairing was reduced. This rationale was supported by our investigation into different MSA pairing strategies; comparing taxonomy-based pairing against simple concatenation revealed that additional pairing did not clearly yield better downstream predictions, especially at higher ipTM threshold (ipTM > 0.8). Thus, we opted for a simple concatenation strategy, which allows the direct re-use of homodimer MSAs, reducing computational demands (**Supplementary Fig. 11**).

#### Structure prediction

Protein complex 3D structure prediction from MSAs was executed either through ColabFold’s *colabfold_batch* command (primarily, for homodimers), or by using an accelerated implementation of OpenFold that leveraged NVIDIA TensorRT and cuEquivariance libraries. Both tools utilised the same set of parameters, namely: one set of weights from AlphaFold-Multimer (model_1_multimer_v3), four recycles/iterations with early stopping, and no relaxation. The outputs from the two inference tools were verified to yield equivalent results, as previously reported^30^. Given the scale of the prediction campaign, the choice of inference parameters was aimed at the reduction of computational demand, while maintaining prediction accuracy. To further reduce overhead, we implemented a parameter in ColabFold (--skip-output msa,plots,pae_json) to only output essential files such as the structure (.pdb) and quality (.json) metrics, and optimised MSA read-in. Failures like out of memory errors or errors due to missing atoms were observed during structure prediction in the presence of large MSAs/long sequences, and non-standard amino acids (e.g. “X”), respectively. These failure modes occurred in a small fraction of the data (<5%), which were not further investigated. Additionally, computation for any one complex was limited to four hours of execution time, and samples that did not produce a structure in the allotted time were discarded.

#### High-performance-compute scaling

In order to increase throughput at scale (i.e., in a multi-GPU, multi-node setting), we wrapped MSA execution and structure predictor in Slurm pipelines which spawned multiple inference processes per GPU on each node, maximising GPU utilisation. GPUs independently processed chunks of data reading from a queueing system, making the inference pipeline naively parallelisable to the number of available GPUs and nodes. The pipelines were executed on a DGX H100 superpod instance, using units of nodes for scale (i.e., at minimum, 8 H100 GPUs).

For ColabFold-based homodimer inference, higher throughput was obtained by packing homodimers of equal length into a single batch to process, sorted by their MSA-depth in descending order. This reduced the amount of Jax (0.7.2) model recompilations, thus increasing prediction throughput. This trick however does not work when processing heterodimers, given that the length of individual chains differs.

For OpenFold, whether for homodimers or heterodimers, this packing strategy is not needed, as the method does not require model re-compilation. However, given a dependency between sequence length and execution time, reserving longer sequences for individual jobs may be beneficial if operating with specific Slurm runtimes. To further accelerate structure prediction, CPU-bound input featurisations were performed for the next input query alongside the GPU-bound inference step for the current query, parallelising heterogeneous computational steps for maximal efficiency.

### Post-processing toolkit and data products

ColabFold and OpenFold outputs (i.e., .pdb and .json files) were converted into AFDB-compliant data products using an extension of the AFDB-Integration-Kit (manuscript in preparation), a multi-stage post-processing toolkit that computes interface and structural quality metrics, generates ModelCIF-compliant mmCIF and Binary CIF files, annotates secondary structure, validates outputs against AFDB schemas, and packages metadata into batched JSON files for database ingestion.

Interface quality metrics such as ipSAE, ipTM, pDockQ2, and LIS were derived from PAE matrices and 3D coordinates, with ipSAE computed through a C++ implementation that processes shards of data in batch mode, increasing throughput. As ipSAE is directional, we computed scores for both chain orderings and defined minimum and maximum such as ipSAE_min_ = min(ipSAE_A→B_, ipSAE_B→A_), to reduce each complex to a single conservative estimate of interface quality. Analogous minimum and maximum operations were applied to LIS and pDockQ2. Backbone and heavy-atom clash counts, and interface residue assignments were identified via GPU-accelerated scripts that pack batches of up to 512 structures into GPU-resident tensors and dispatch atomic distance kernels with torch_cluster.radius_graph. Secondary structure annotation was inferred with PyDSSP (github.com/ShintaroMinami/PyDSSP; commit e251a43), a vectorised NumPy/PyTorch implementation that eliminates subprocess overhead and processes entire batches in memory. JSON parsing in the toolkit was replaced with *orjson* (3.11.4), reducing deserialisation of large PAE matrices by roughly an order of magnitude. Similarly, PDB structure parsing was replaced with *Gemmi*^44^.

Each post-processing stage parallelises its own work across ‘ProcessPoolExecutor’ process pools, bypassing the Python GIL for CPU-bound operations, while metadata lookups were batched into a single DuckDB query per data shard, and cached in module-level globals shared across workers, eliminating per-model database roundtrips. DuckDB memory usage is capped to prevent out-of-memory events on shared nodes, and downstream shard aggregation streams data through PyArrow batches to avoid loading full datasets into memory.

To distribute this workload across nodes in our HPC setting, we implemented a Slurm array job framework based entirely on file-level coordination: a driver script partitions a flat list of model IDs into contiguous shards of at most 5,000 structures per array task, with each task operating independently and deterministically from its array index alone. Each node writes outputs to local scratch memory, uploads completed batches directly to object storage via ‘s5cmd’, and records only small marker files on a shared Lustre filesystem, avoiding I/O contention at scale. Fault tolerance was achieved through two-level automatic resume, skipping completed pipeline stages within a shard and exiting immediately for already-processed shards on resubmission, making runs across hundreds of nodes robust to node failures or Slurm job preemption.

### Benchmark dataset construction

We constructed precision-focused, non-redundant benchmark sets by integrating SIFTS-based^45^ AFDB-to-PDB mappings with Foldseek clustering, full-PDB decontamination, coverage filtering, and post-training exclusion.

#### Input processing and categorisation

Starting from 3,867,387 SIFTS-derived AFDB-to-PDB chain mappings (106,411 predicted structures, 172,988 PDB entries), we retained X-ray crystallography, cryo-EM, and neutron diffraction structures with author-defined biological assemblies (3,705,034 mappings). Each PDB entry was categorised by stoichiometry as homodimer, monomer, heterodimer, or hetero-*N*-mer. When a predicted structure mapped to PDB entries with conflicting stoichiometries, evidence for an interacting assembly was prioritised, and the structure was excluded from the corresponding negative pool.

#### Clustering and redundancy removal

Structural redundancy was removed using Foldseek-Multimer commit 95b9adb. Dimers were clustered with easy-multimercluster using -c 0.6, --chain-tm-threshold 0.7, --interface-lddt-threshold 0.3, and --cov-mode 0, yielding 15,534 homodimer and 14,325 heterodimer clusters. Monomeric chains were clustered with *easy-cluster* using -c 0.6, --tmscore-threshold 0.7, and --cov-mode 0, yielding 4,696 monomer and 24,112 hetero-N-mer chain clusters. Cluster representatives were selected by total modelled chain sequence length, preferentially retaining full-length structures over fragmentary ones.

For negative sets, entries were excluded when none of the mapped PDB entries covered at least 70% of the corresponding UniProt sequence, because partial-domain structures were insufficient to support a negative label for the corresponding full-length predicted entry. In addition, PDB entries annotated as truncations or artificial constructs in mmCIF metadata were excluded from negative-source categories (167 monomer and 62 hetero-*N*-mer entries). Positive sets were not coverage-filtered, because even a partial experimental structure can provide direct evidence for an observed oligomeric state or physical interaction.

#### Post-training filtering

To ensure independence from the AlphaFold-Multimer training set, structures were searched against the full PDB using Foldseek *easy-multimersearch* for dimers and *easy-search* for monomers. For dimer-level searches, a PDB-derived dimer database was generated using *createdimerdb* in Foldseek-Multimer commit 95b9adb. Predicted dimers were first searched against this database with the following parameters: --cov-mode 0 --interface-lddt-threshold 0.1 --chain-tm-threshold 0.1 --tmscore-threshold 0. Hits were then post-filtered and retained only if they satisfied all of the following criteria: either if both query-chain TM-scores were at least 0.7 or both target-chain TM-scores were at least 0.7, the interface lDDT score was at least 0.3, and either query or target coverage was at least 0.6. For monomer-level searches, hits were retained if coverage was ≥ 0.6 and either query or target TM-score ≥ 0.7.

For the homodimer benchmark (both positive and negative) and the heterodimer positive set, a cluster representative was retained only if every member of its cluster had a post-training PDB release date and no structural homologs in pre-training PDB entries; a single pre-training match or missing search result within a cluster excluded the entire group. For the heterodimer negative set, only post-training PDB release dates for all cluster members were required, as the non-interaction label derives by the absence of physical contact between the two entities within experimentally observed assemblies rather than the novelty of individual subunits.

#### Homodimer benchmark

Homodimer positives were derived from 5,551 post-training PDB entries annotated as experimentally supported homodimeric assemblies. After Foldseek clustering, cluster-level post-training filtering, SIFTS-based mapping to AFDB accessions, this yielded 230 non-redundant AFDB accessions.

Homodimer negatives were derived from 5,217 post-training PDB entries annotated as monomeric assemblies. Candidate monomer chains were screened against the full PDB to remove queries with homo-oligomeric Foldseek structural homologs, retaining monomer-level hits at alignment coverage ≥ 0.6 and max(qTM, tTM) ≥ 0.7. After Foldseek monomer-cluster representative selection, coverage filtering, cluster-level post-training filtering, and SIFTS-based mapping to AFDB accessions, this yielded 117 non-redundant AFDB accessions.

#### Heterodimer benchmark

Heterodimer positives were derived from experimentally supported heterodimeric PDB assemblies in which both subunits mapped to the same predicted AFDB heterodimeric pair. This identified 4,525 PDB chain pairs from 3,560 post-training PDB entries. After Foldseek heterodimer clustering, 1,792 clusters had at least one member with both chains traceable to an AFDB pair through SIFTS, from which one representative was retained per cluster. Cluster-level post-training filtering and AFDB pair mapping yielded 94 non-redundant AFDB pair IDs.

Heterodimer negatives were constructed from predicted heterodimer pairs whose subunits co-occurred in the same hetero-N-mer PDB assembly but did not physically interact in any PDB entry containing both entities. Interaction status was assessed at the entity-pair level across all PDB entries containing both entities, using a polymer chain mapping table to bridge SIFTS label_asym_id and Foldseek auth_asym_id identifiers. After retaining at most one AFDB pair per Foldseek cluster pair, applying cluster-level post-training and coverage filters, and verifying the absence of Foldseek interactions in any PDB entry, this yielded 250 non-redundant AFDB heterodimeric pairs.

### Analysis

#### Predicted complex structures for downstream analyses

Of the 1,735,475 high-confidence homodimeric and 79,392 heterodimeric entries, 35 homodimer structure files were unavailable at the time of analysis; all downstream structure-based analyses therefore used 1,735,440 homodimers and 79,392 heterodimers.

#### Structural coverage analysis

To quantify the extent to which predicted complexes expand structural coverage beyond experimentally determined multimers, we compared the number of high-confidence predicted complexes (1,735,440 homodimers and 79,392 heterodimers) with the number of experimentally determined multimeric structures deposited in the Protein Data Bank that are mapped to UniProt (downloaded in March 2026), per organism. PDB multimer counts were tallied by NCBI Taxonomy^46^ identifier, and taxonomic lineage information was resolved using NCBI taxonomy files (names.dmp, nodes.dmp). Organisms with fewer than 10 high-confidence predicted structures were excluded to avoid unreliable estimates. For organism-level comparison, each species was represented by both its PDB multimer count and its predicted high-confidence complex count. For kingdom-level comparison, taxa were aggregated into seven broad categories: Metazoa, Viridiplantae, Fungi, Other Eukaryotes, Bacteria, Archaea, and Viruses. Organisms with no corresponding PDB multimer entry were retained and displayed with predicted counts only.

#### Sequence cluster comparison

Sequence clusters were generated using MMseqs2^37^ (release 18) from the 1,735,440 high-confidence homodimer chain sequences. Cluster representatives were obtained by clustering sequences at 98% sequence identity and 95% coverage using the parameters --min-seq-id 0.98 -c 0.95 --cov-mode 0. Structural similarity within each cluster was evaluated using Foldseek^39^ (commit d6204679). A Foldseek database was constructed from the 1,735,440 high-confidence structures, and representative structures were aligned against cluster members using foldseek *structurealign* with --alignment-type 2. Structural similarity between representative and member structures was quantified using TM-score.

#### Within-homodimer chain comparison

To assess structural consistency between chains in predicted homodimers, 1,735,440 structural alignments were performed between chain A and chain B of each complex using Foldseek (commit d6204679). A Foldseek database was constructed from the 1,735,440 high-confidence structures, and chain pairs were aligned using monomer structural alignment. A prefilter mapping was used to directly match chain A to chain B for each predicted complex, and structural similarity was quantified using Foldseek TM-score metrics.

#### Heterodimer physical-contact enrichment analysis

To assess whether high-confidence heterodimer predictions are enriched among experimentally observed direct-contact pairs, we compiled a separate post-training set of 617 STRING-derived AF heterodimer pairs. This set used the same entity-pair interaction classification, chain-identifier bridging and post-training filtering as the heterodimer benchmark, but also included interacting pairs from hetero-oligomeric PDB complexes and applied cluster-pair deduplication jointly across heterodimer and hetero-N-mer cluster spaces. The final set contained 371 physically interacting pairs and 246 non-interacting controls.

#### Mapping to AFDB v6 monomeric models

For monomer-dimer pLDDT comparisons, chains from our UniProt 2025_04 complex predictions were mapped to AFDB v6 monomeric models, which were based on UniProt 2025_03. To avoid release-version mismatches, we excluded AFDB entries listed as added, removed or changed in the AFDB *diffs.ndjson.gz* file and retained only unchanged entries mapped by UniProt accession. This yielded 12.8 million sequence-matched homodimer predictions and 7,520,086 heterodimer predictions for which both chains had matched AFDB v6 monomeric models.

#### Clustering analyses

1,814,832 structures from 1,735,440 high-confidence homomeric and 79,392 high-confidence heterodimeric structures were clustered using Foldseek Multimercluster (commit d6204679). Cluster representatives were obtained by clustering structures at 60% 3Di sequence coverage, 30% interface similarity and 70% chain similarity using the parameters -c 0.6 --interface-lddt-threshold 0.3 --chain-tm-threshold 0.7 --cov-mode 0. These parameters follow those used for the Teddymer dataset^47^.

#### Taxonomic analysis

Taxonomic lineages and rank assignments were resolved using the NCBI Taxonomy database (taxdump, downloaded in February 2026); specifically, nodes.dmp and names.dmp were used to construct parent–child relationships and to assign each taxon to standardised ranks (e.g., superkingdom, phylum). These rank assignments were used both for computing cluster-level lowest common ancestors (LCAs) and for stratifying prediction success rates across taxonomic clades (**Fig. 3a,b)**. Based on UniProt taxonomic assignments and using the MMseqs2 taxonomy’s lca module^48^, we computed, for each non-singleton cluster, the LCA from its member taxa, and visualised the resulting taxonomic distribution as a Sankey diagram using Metabuli-App^49^.

#### Structural search of the high-confidence subset against PDB

To assess structural coverage against experimentally determined structures, we searched 1,735,440 high-confidence homomeric and 79,392 high-confidence heterodimeric predicted structures against the PDB, downloaded in January 2026, using Foldseek-Multimer (commit d6204679). A dimer database was first generated from the PDB using Foldseek’s *createdimerdb* module (commit 95b9adb). The 1,814,832 predicted dimers were then searched against this database with the following parameters: --cov-mode 0 --interface-lddt-threshold 0.1 --chain-tm-threshold 0.1 --tmscore-threshold 0. Hits were then post-filtered and retained only if they satisfied all of the following criteria: either if both query-chain TM-scores were at least 0.7 or both target-chain TM-scores were at least 0.7, the interface lDDT score was at least 0.3, and either query or target coverage was at least 0.6. These thresholds were selected based on the Teddymer dataset. Because many PDB structures are truncated, the final filtering was performed manually rather than relying solely on Foldseek input parameters.

## Supplementary Material

**Supplementary Figure 1.**
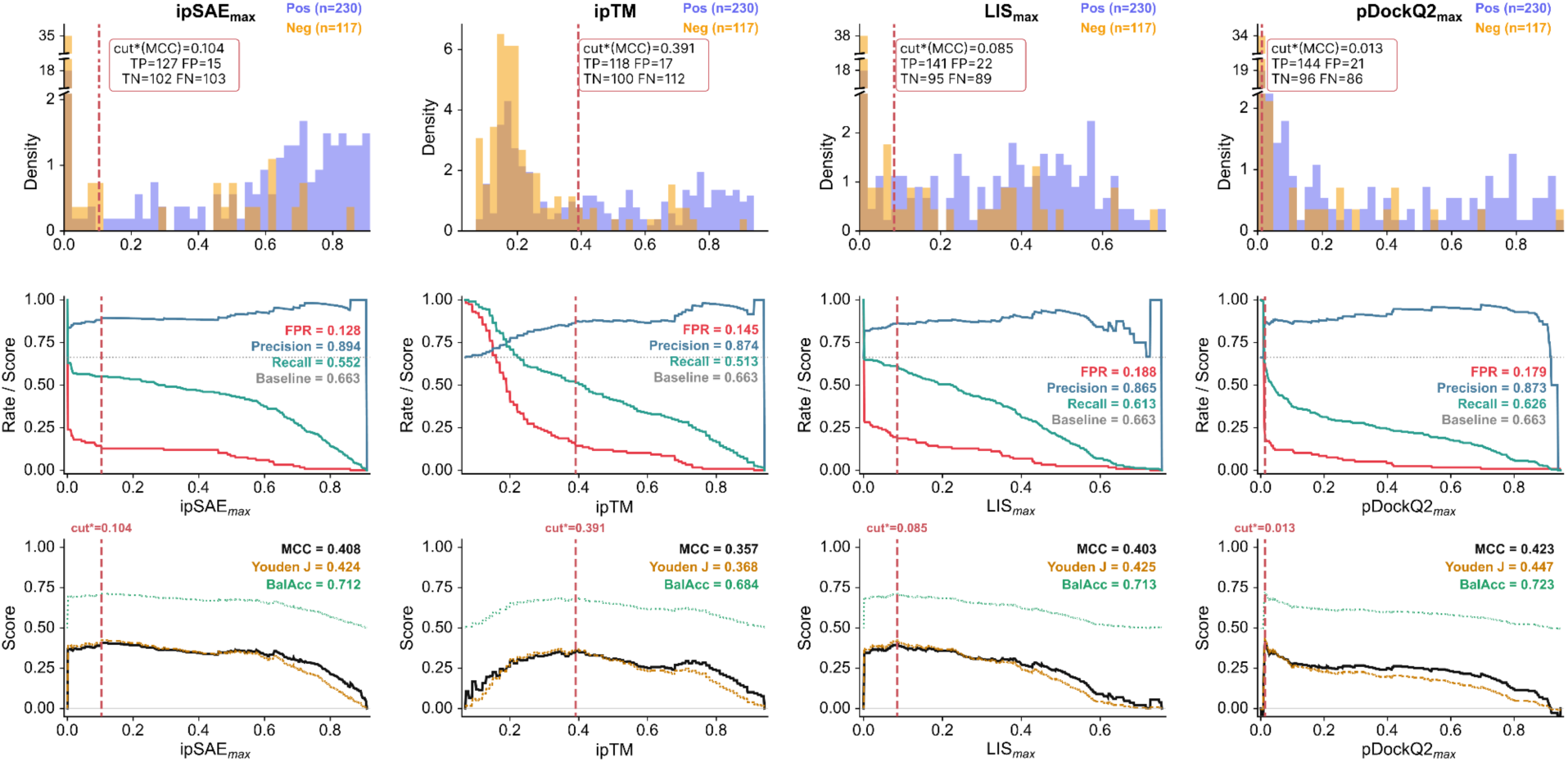
Binary classification performance of four confidence metrics on the homodimer post-training benchmark set, with cutoffs selected by argmax (MCC). **a,** ipSAE_max_ (cut*=0.104), **b,** ipTM (cut*=0.391), **c,** LIS_max_ (cut*=0.085), **d,** pDockQ2_max_ (cut*=0.013). Top row: score distributions of positives (blue, n=230) and negatives (orange, n=117) with confusion matrix counts at the MCC-optimal cutoff (dashed dark red line). Middle row: FPR, precision and recall versus threshold. Bottom row: MCC, Youden’s J and balanced accuracy versus threshold. Post-training subset restricted to structures whose PDB structural homologs were released after 2021-09-30.

**Supplementary Figure 2.**
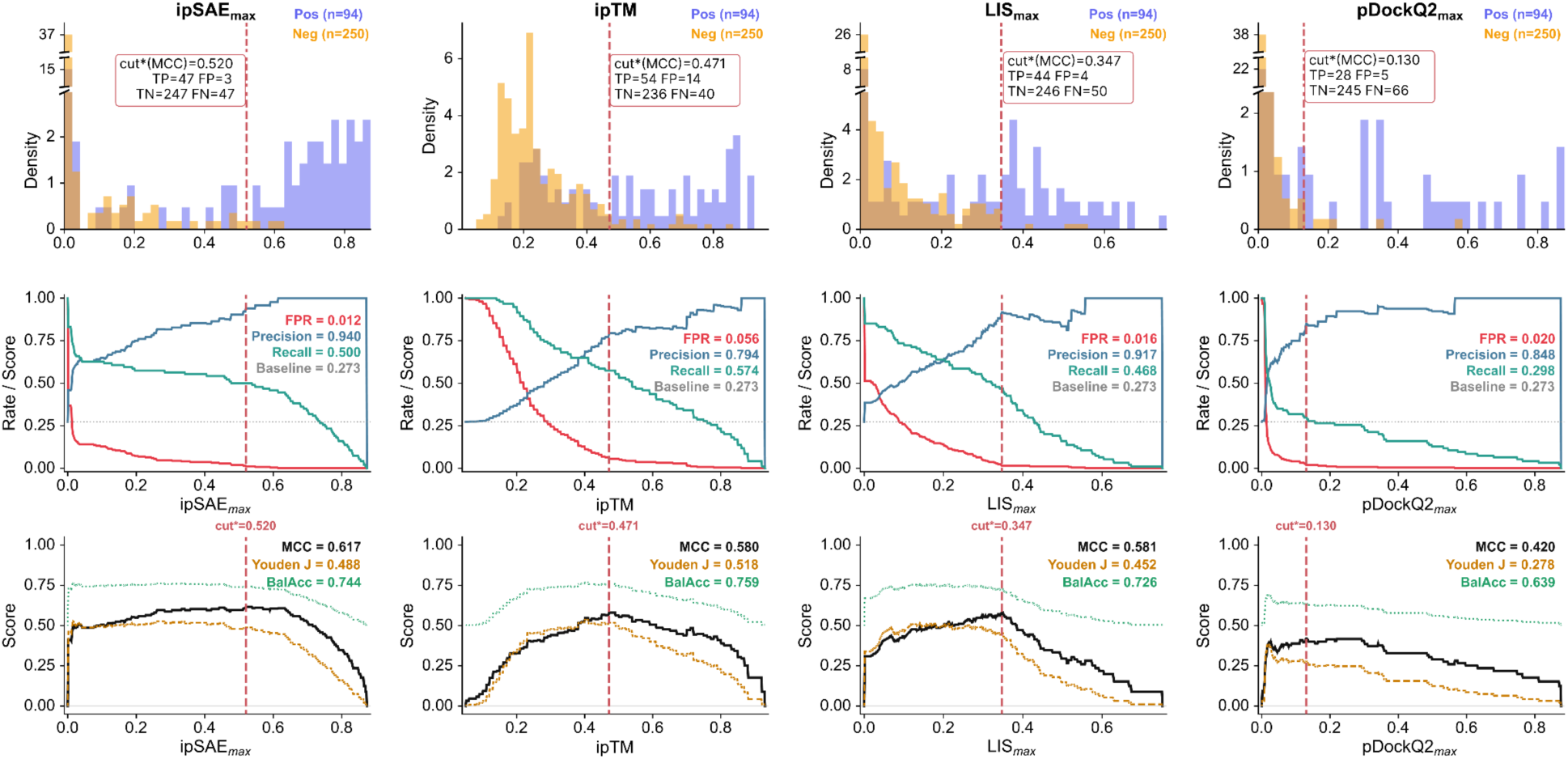
Binary classification performance of four confidence metrics on the heterodimer post-training benchmark set, with cutoffs selected by argmax (MCC). **a,** ipSAE_max_ (cut*=0.520), **b,** ipTM (cut*=0.471), **c,** LIS_max_ (cut*=0.347), **d,** pDockQ2_max_ (cut*=0.130). Top row: score distributions of positives (blue, n=94) and negatives (orange, n=250) with confusion matrix counts at the MCC-optimal cutoff (dashed red line). Middle row: FPR, precision and recall versus threshold. Bottom row: MCC, Youden’s J and balanced accuracy versus threshold. Post-training subset restricted to structures whose PDB structural homologs were released after 2021-09-30.

**Supplementary Figure 3:**
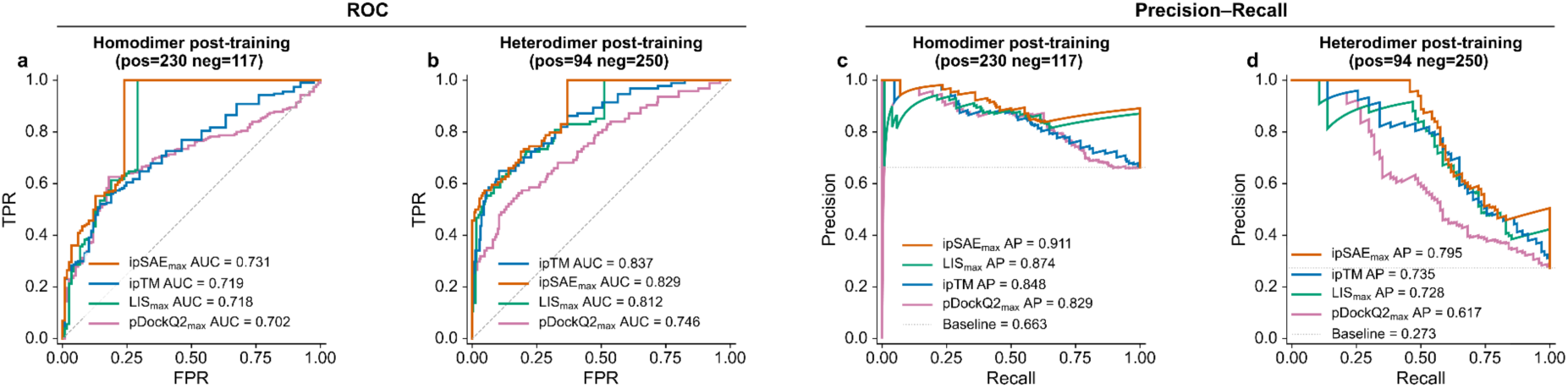
Receiver Operating Characteristic (ROC) and Precision–Recall curves for four multimer prediction confidence metrics on post-training benchmark sets. **a,** Homodimer ROC (pos=230, neg=117). **b,** Heterodimer ROC (pos=94, neg=250). **c,** Homodimer PR. **d,** Heterodimer PR. (e–h) Same as a–d but for Precision–Recall. Metrics: ipSAE_max_ (vermillion), ipTM (blue), LIS_max_(bluish green), pDockQ2_max_ (reddish purple). Legend entries sorted by descending AUC or AP. Dashed diagonal (ROC) and dotted line (PR, at prevalence) indicate chance level.

**Supplementary Figure 4:**
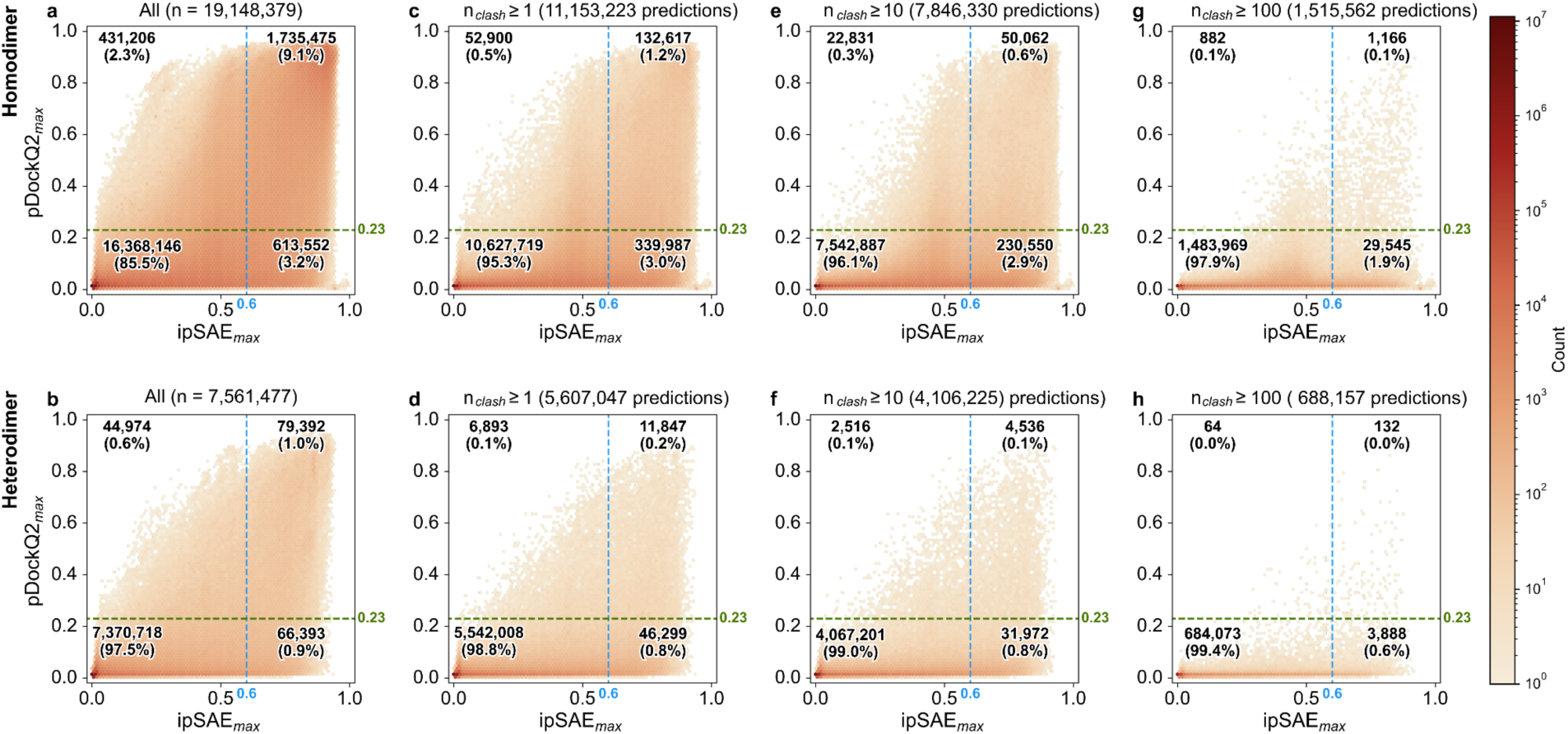
Backbone-clash distributions under joint ipSAE_max_ and pDockQ2_max_ filtering. Hexbin density plots of ipSAE_max_ versus pDockQ2_max_ for homodimer (top row) and heterodimer (bottom row) AlphaFold-Multimer predictions, stratified by backbone clash severity. **a,b,** All structures. **c,d,** Structures with at least one backbone clash (n_clash_ ≥ 1). **e,f,** n_backbone_clash_ ≥ 10. **g,h,** n_clash_ ≥ 100. Dashed lines indicate the fixed cutoffs (ipSAE_max_ =0.6, blue; pDockQ2_max_ =0.23, orange). Quadrant counts and percentages shown in each panel. Shared log-scale colorbar. Homodimer: 19,148,379 predictions; heterodimer: 7,561,477 predictions.

**Supplementary Figure 5:**
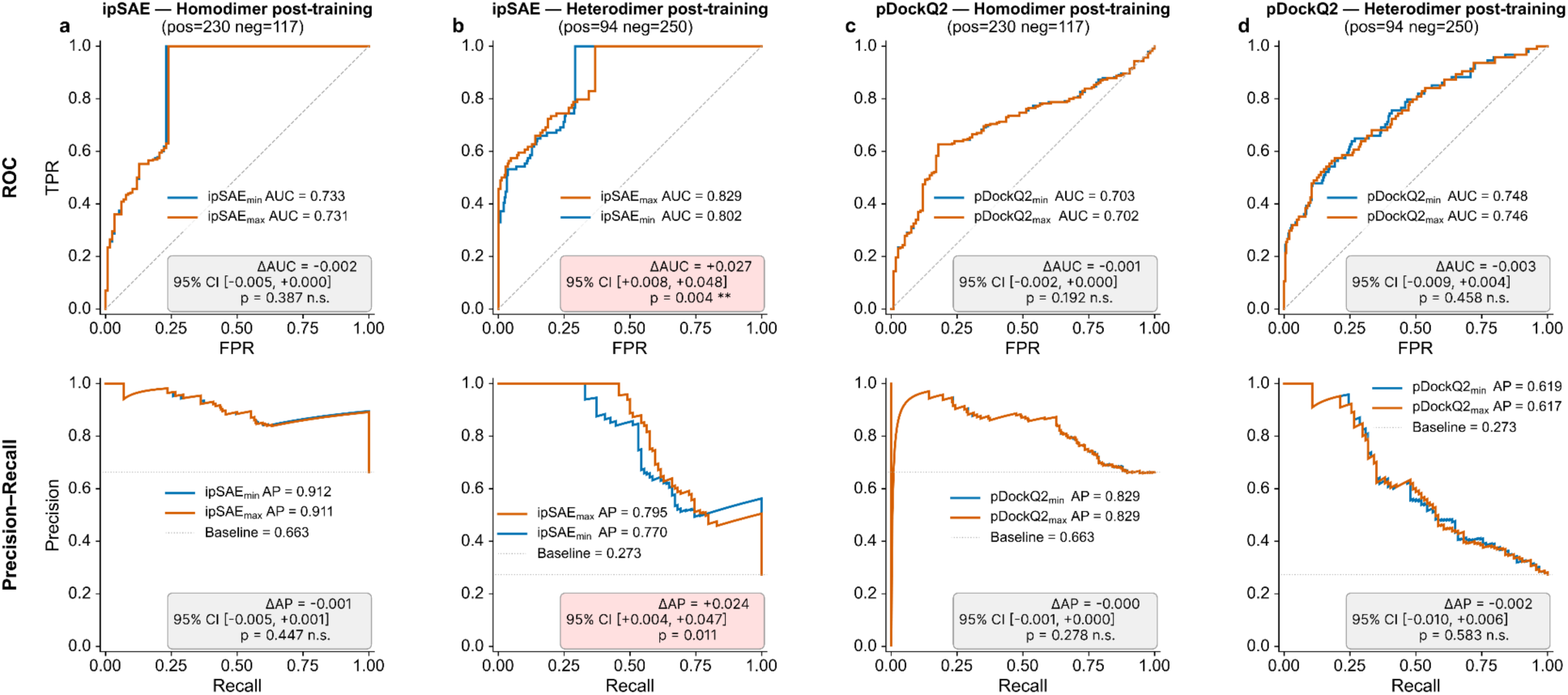
Comparison of max- and min- chain-level score aggregation of ipSAE and pDockQ2 on post-training benchmark sets. **a,** ipSAE, homodimer (pos=230, neg=117). **b,** ipSAE, heterodimer (pos=94, neg=250). **c,** pDockQ2, homodimer. **d,** pDockQ2, heterodimer. Top row: ROC curves. Bottom row: Precision–Recall curves. Each panel shows a paired bootstrap test (2,000 iterations): delta, 95% CI, and two-sided p-value. Pink background indicates statistical significance (p<0.05); grey indicates non-significance. Max (vermillion), min (blue).

**Supplementary Figure 6:**
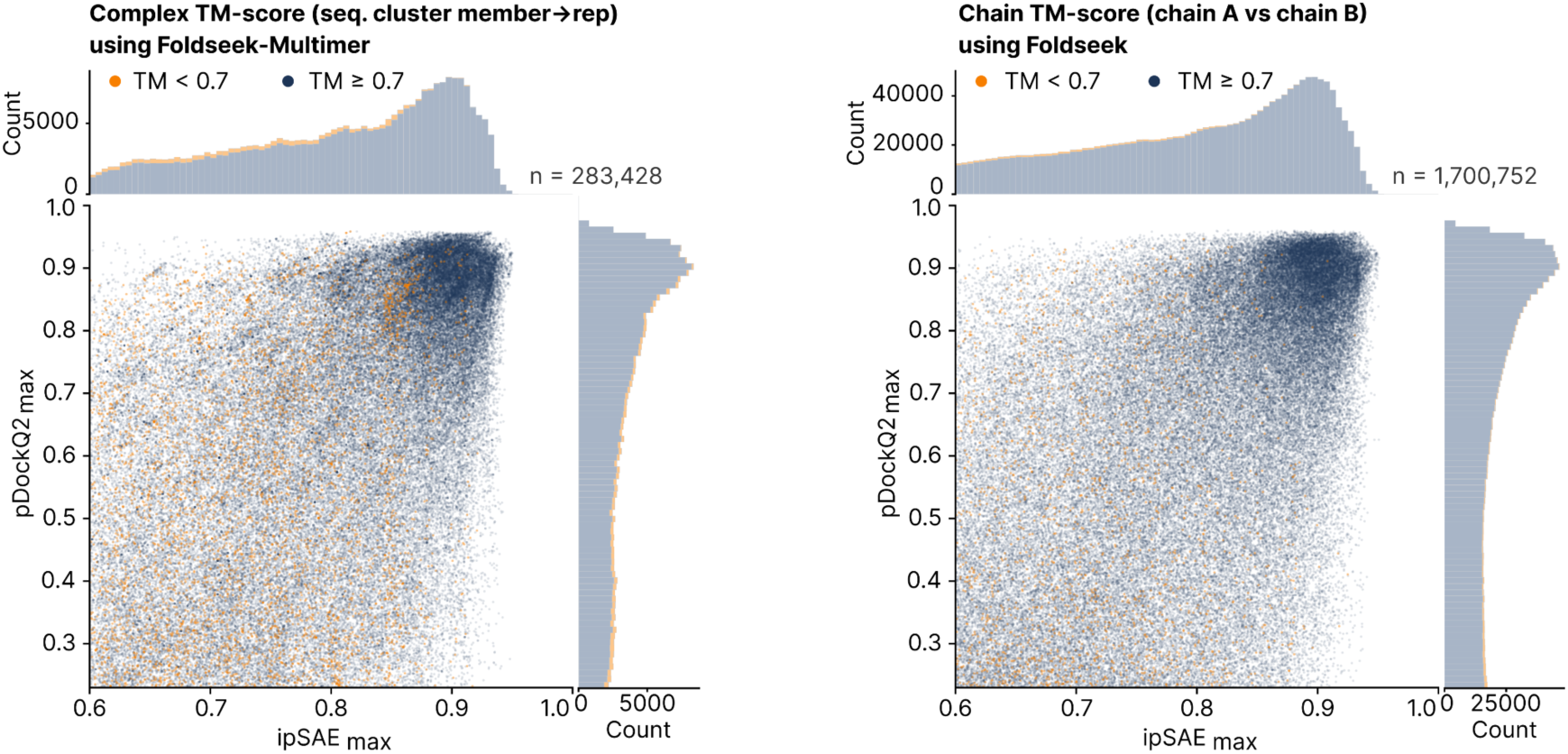
Structural consistency of high-confidence homodimers across sequence-similar clusters and between chains. **a,** ipSAE_max_ versus pDockQ2_max_ for each high-confidence (ipSAE_max_ ≥ 0.60, pDockQ2_max_ ≥ 0.23) homodimers (n = 283,428 members of MMseqs2 98%-identity, 95%-coverage non-singleton clusters), coloured by the complex TM-score (max(qtm, ttm), Foldseek-Multimer) to the non-self cluster representative: 267,990 (94.55%) reach TM ≥ 0.7 (navy), the remainder fall below (orange). **b,** Same axes for all high-confidence homodimer predictions with max_tm_AB defined (n = 1,700,752), coloured by the chain-A vs chain-B TM-score within each prediction (max_tm_AB, Foldseek): 1,664,813 (97.89%) reach TM ≥ 0.7 (navy), the remainder fall below (orange). Marginal histograms (TM ≥ 0.7 slate at base, TM < 0.7 peach on top) use the full populations; scatter shows 100,000 random points per panel (random_state = 42).

**Supplementary Figure 7.**
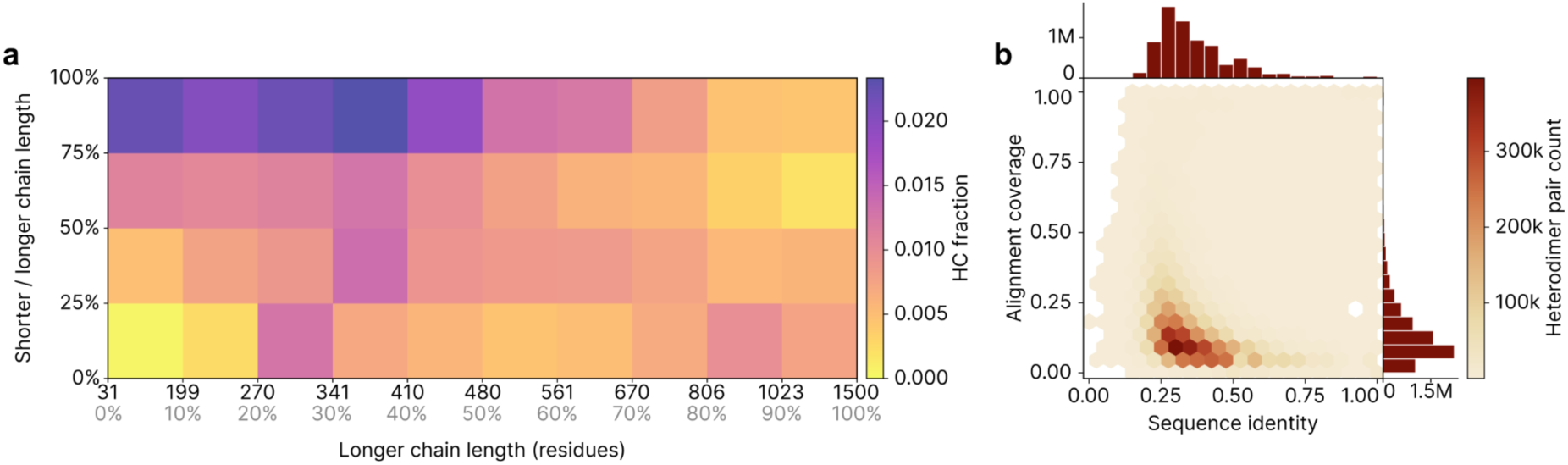
Heterodimer high-confidence fractions vary with chain length symmetry and sequence similarity. **a,** Fraction of high-confidence predictions across STRING-derived heterodimer candidate pairs, stratified by longer-chain length and shorter-to-longer chain-length ratio. Longer-chain length was divided into deciles, and chain-length ratio into quartiles. High-confidence predictions were defined as ipSAE_max_ ≥ 0.60 and pDockQ2_max_ ≥ 0.23; the dotted line on the colour scale marks the overall high-confidence fraction of 1.05%. High-confidence predictions are enriched among mid-length, length-symmetric pairs. **b,** Hexbin density of pairwise sequence identity versus alignment coverage between the two chains, with marginal histograms. The high-identity, high-coverage region, defined as sequence identity ≥ 0.90 and alignment coverage ≥ 0.80, represents a homodimer-like subset with a higher high-confidence fraction than the full heterodimer set (15.1% versus 1.05%), but contains only 6,418 pairs, corresponding to 0.085% of all heterodimer candidates.

**Supplementary Figure 8.**
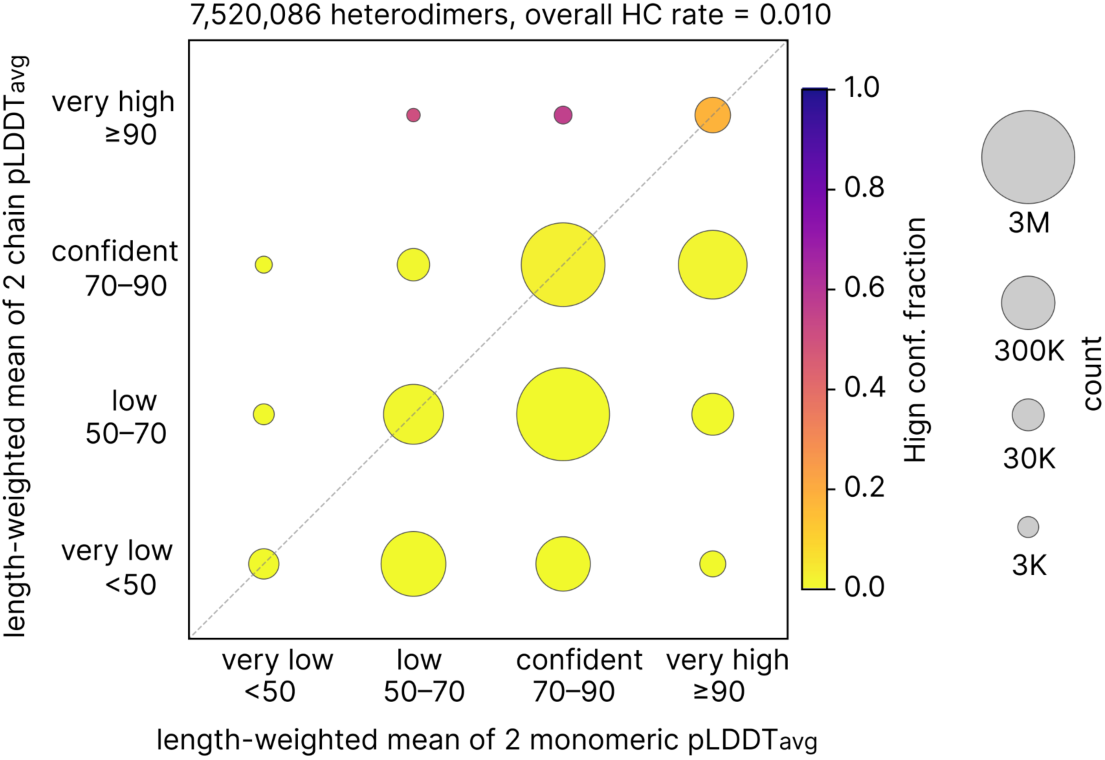
Heterodimer confidence across monomer-dimer pLDDT regimes. STRING-derived heterodimer predictions for which both chains could be mapped to unchanged AFDB v6 monomeric models were binned by length-weighted mean monomer pLDDT and heterodimer pLDDT (n = 7,520,086). Bins correspond to AFDB confidence intervals (<50, 50-70, 70-90, ≥90). Bubble area is proportional to the square root of pair count, and colour indicates the within-cell fraction passing the high-confidence criterion, ipSAE_max_ ≥ 0.60 and pDockQ2_max_ ≥ 0.23. The dashed line marks equal monomer and dimer pLDDT. Most pairs lie on or below the diagonal; collapse-like cells, defined as monomer pLDDT ≥70 and dimer pLDDT <70, contain ∼46% of pairs. High-confidence predictions are enriched where both monomer and dimer pLDDT are very high (n = 49,964; 17.6%) and, to a smaller extent, where monomer pLDDT is confident and dimer pLDDT is very high (n = 912; 55.7%).

**Supplementary Figure 9:**
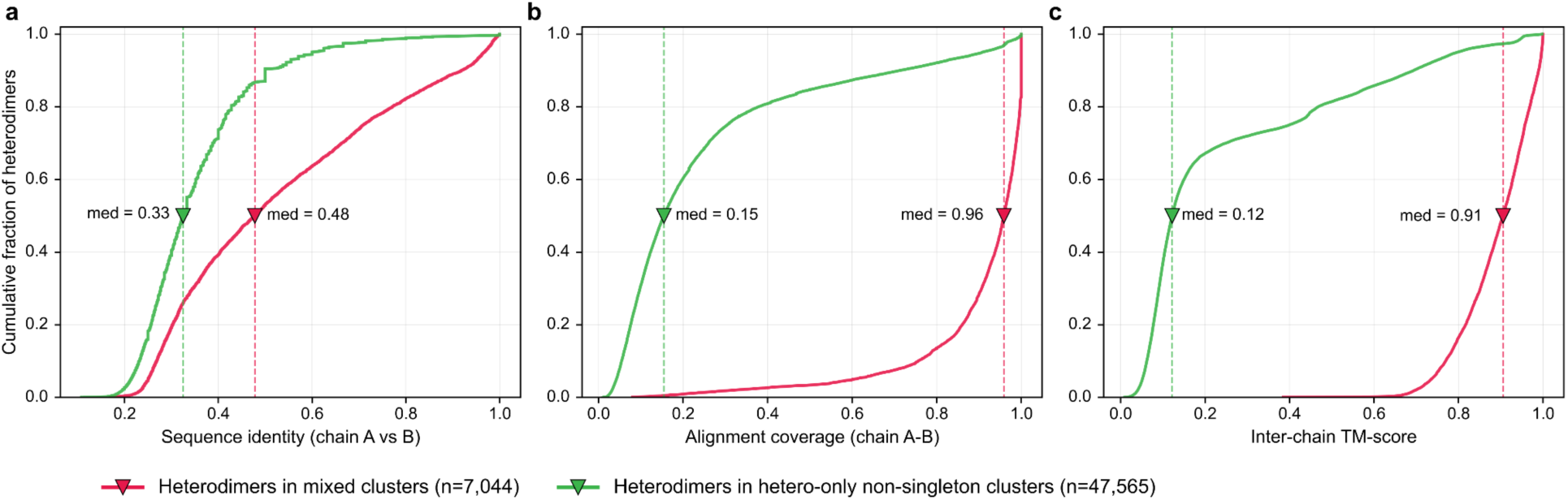
Heterodimers that co-cluster with homodimers are structurally homodimer-like. Cumulative distribution functions of three chain-pair similarity metrics for every heterodimer in the FS-MM clusters, stratified by the composition of the host cluster: red, the 7,044 heterodimers in mixed clusters (1,845 clusters that also contain at least one homodimer); green, the 47,565 heterodimers in 7,356 hetero-only clusters. **a,** sequence identity between the two chains. **b,** best per-chain alignment coverage on the A-B alignment. **c,** best inter-chain TM-score. Triangles and dashed lines mark per-group medians. The red distribution is shifted to the right of the green in every panel (medians 0.48 versus 0.33, 0.96 versus 0.15 and 0.91 versus 0.12), indicating that heterodimers sharing a structural cluster with a homodimer have chains of highly similar fold and near-complete mutual coverage and are predominantly homodimer-like complexes.

**Supplementary Figure 10:**
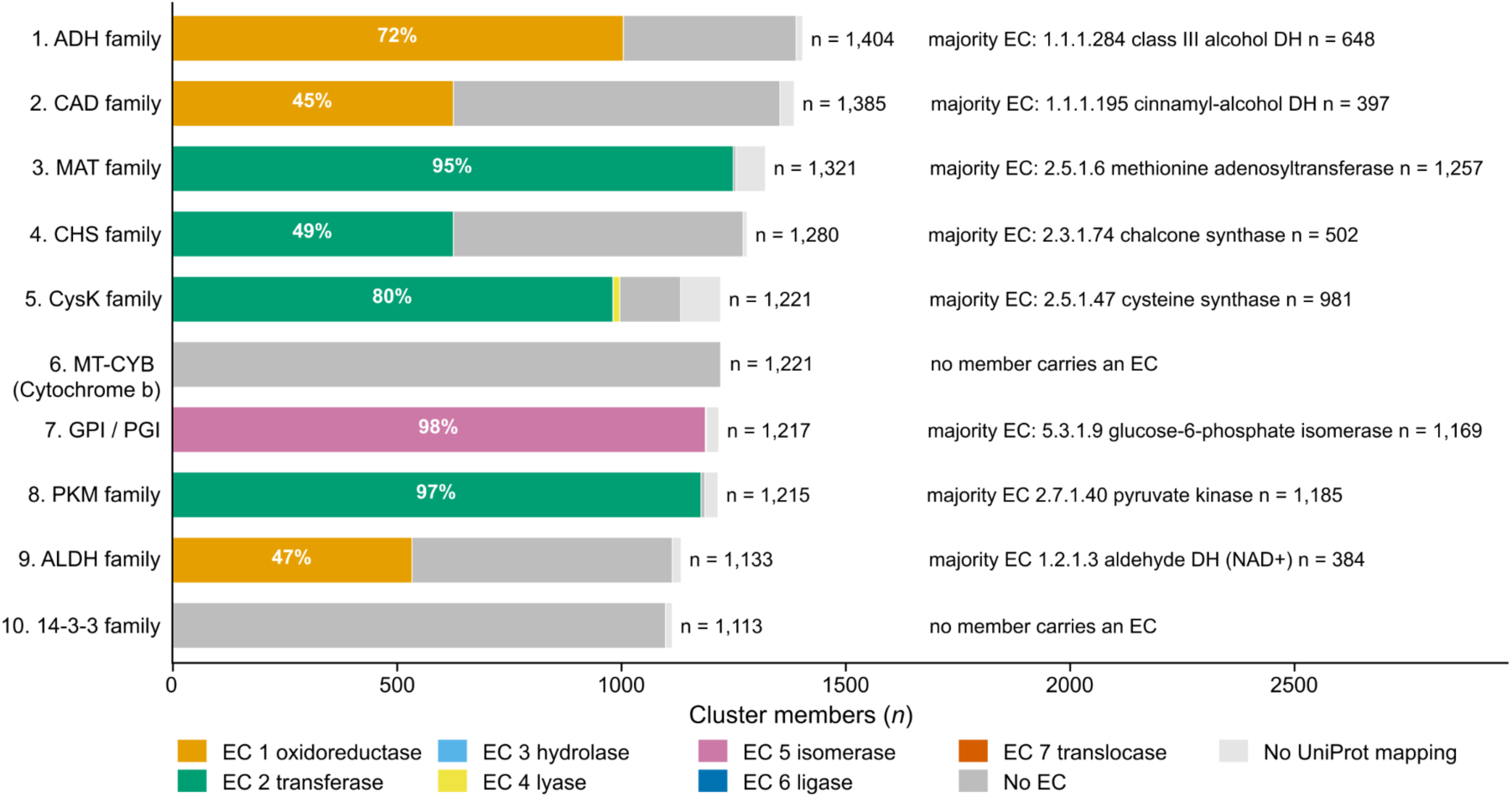
Eight of the ten largest complex-level structural clusters are dominated by a single Enzyme Commission (EC) class; the remaining two carry no EC annotation. Bars (rank 1 top to rank 10 bottom) are partitioned by the UniProt ‘ec’ field into the seven IUBMB EC classes (colour-coded), ‘No EC’ (UniProt entry present, field empty) and ‘No UniProt mapping’. The right-side text gives the total cluster size (first n), the most frequent EC within the cluster (’majority EC’) with its IUBMB recommended name, and the number of members carrying that EC (second n). The two non-EC clusters are mitochondrial cytochrome b (rank 6, a Complex III subunit; IUBMB does not assign EC at the single-subunit level) and the 14-3-3 family (rank 10, a non-catalytic adapter). Because the UniProt ‘ec’ field is sparsely populated for entries that have not been manually curated, the EC counts shown here are a lower bound on the true catalytic content of each cluster.

**Supplementary Figure 11:**
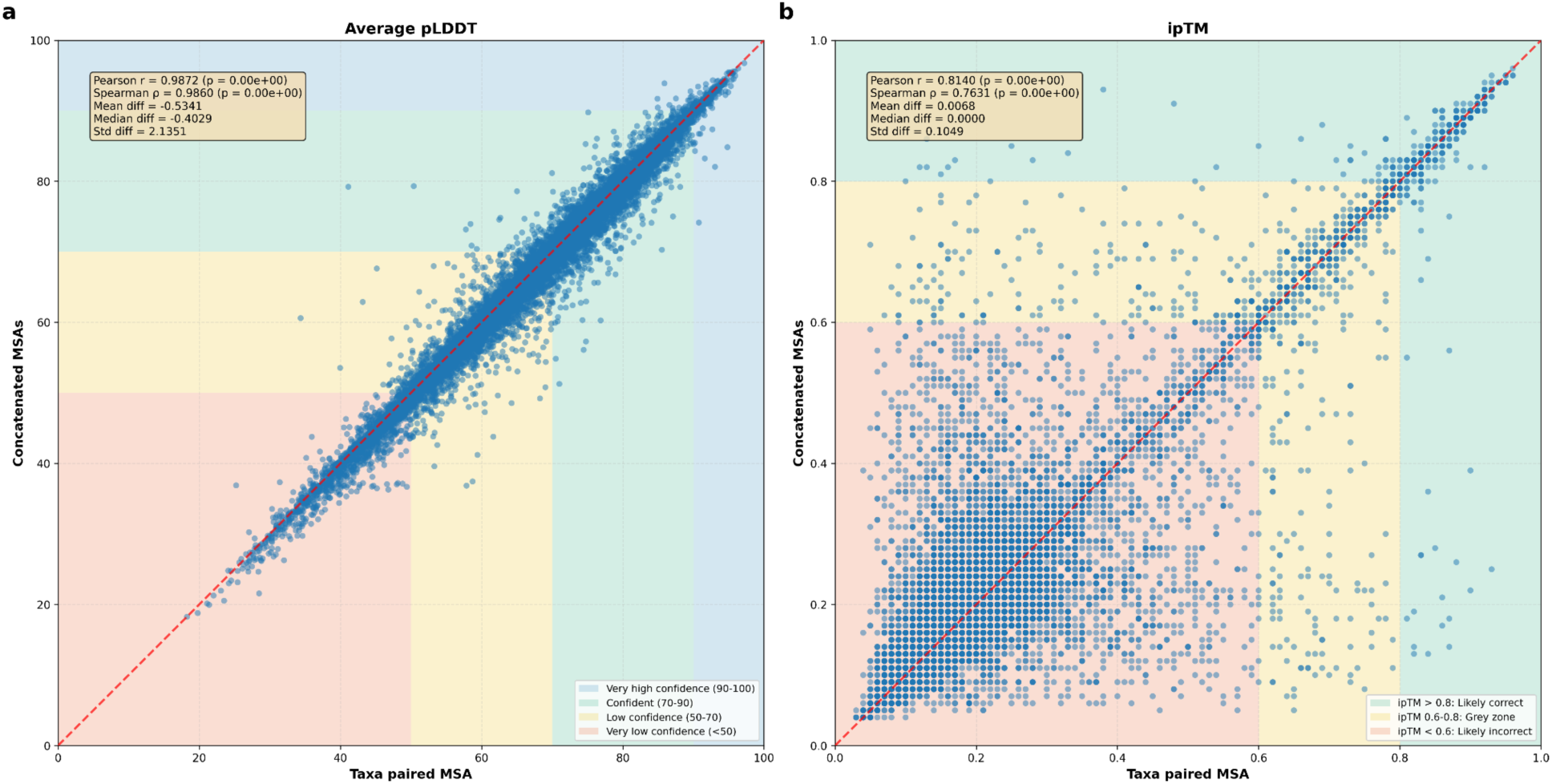
Comparison of pLDDT (a) and ipTM (b) scores for predicted structures using ColabFold (via colabfold_batch) using two distinct MSA generation strategies as inputs. On the x axis: taxa paired MSAs following the standard colabfold_search pipeline with MMseqs2-GPU as a backend. On the y axis: concatenated homodimer (taxa filtered monomer) MSAs computed with colabfold_search using MMseqs2-GPU as a backend. For average pLDDT, there is generally great agreement between the two input MSA strategies (plot coloured by AFDB defined confidence bands). For ipTM, there is also general agreement, with more noise in the “likely incorrect” regions, and narrower limits of agreement with greater correlation in the “likely correct” region, despite outliers in both directions.

**Supplementary Table 1:**
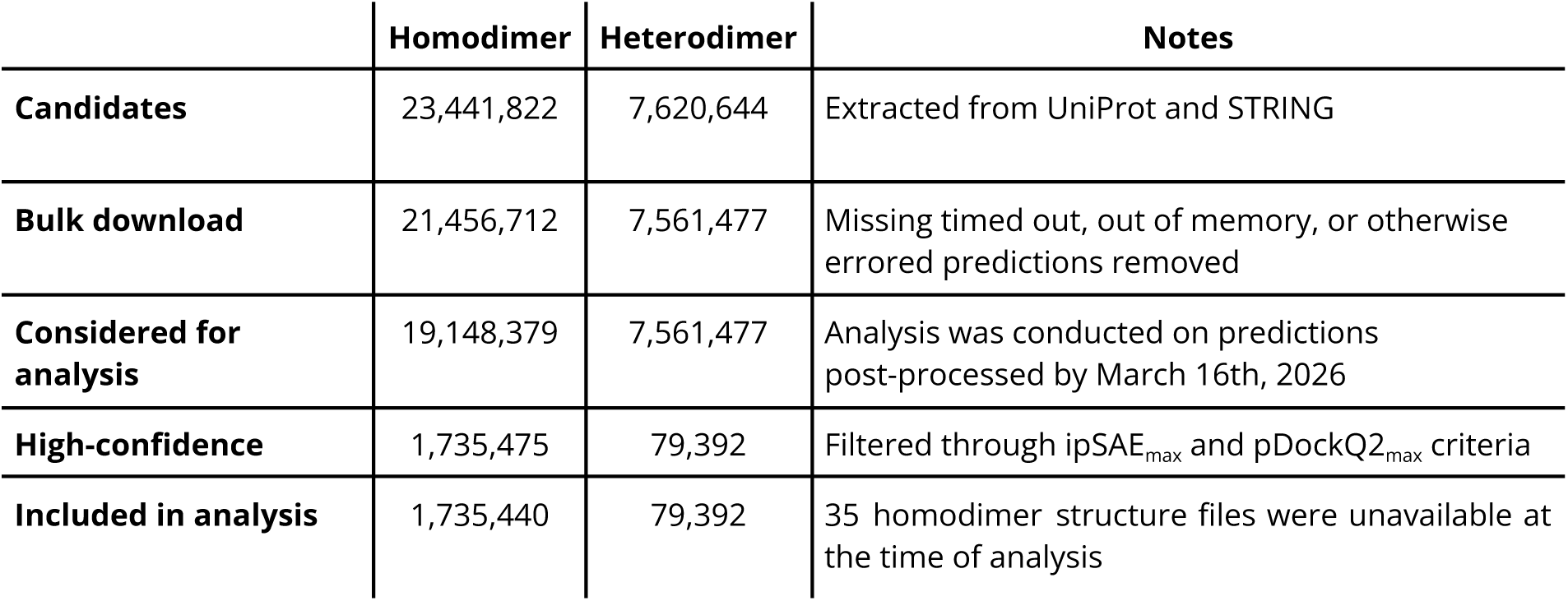
Summary of heterodimer and homodimer dataset filtering and analysis set construction.

|  | Homodimer | Heterodimer | Notes |
| --- | --- | --- | --- |
| Candidates | 23,441,822 | 7,620,644 | Extracted from UniProt and STRING |
| Bulk download | 21,456,712 | 7,561,477 | Missing timed out, out of memory, or otherwise errored predictions removed |
| Considered for analysis | 19,148,379 | 7,561,477 | Analysis was conducted on predictions post-processed by March 16th, 2026 |
| High-confidence | 1,735,475 | 79,392 | Filtered through ipSAE<sub>max</sub> and pDockQ2<sub>max</sub> criteria |
| Included in analysis | 1,735,440 | 79,392 | 35 homodimer structure files were unavailable at the time of analysis |

In the following, all proteomes included in our study (compressed to PT5 for efficiency):

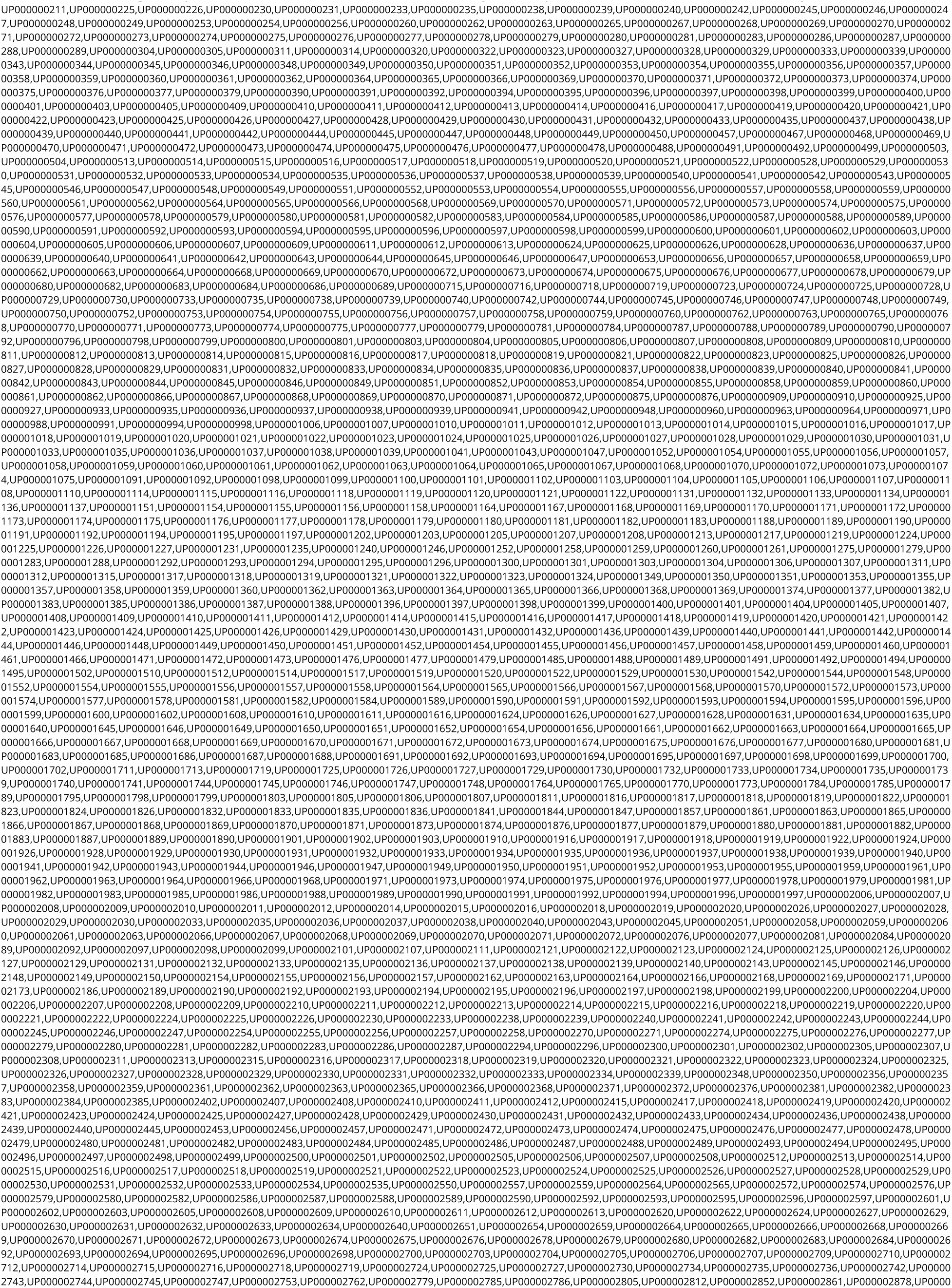
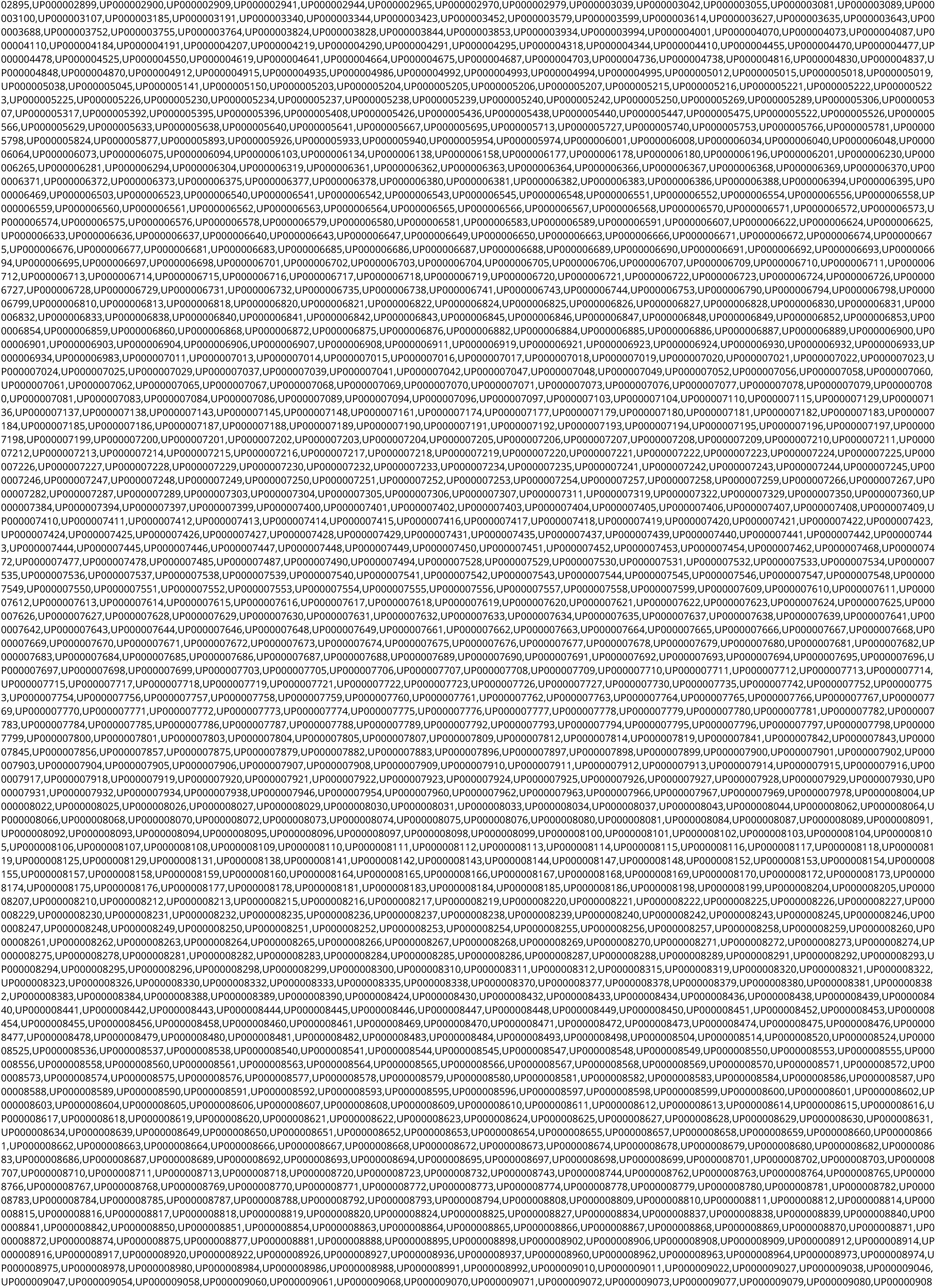
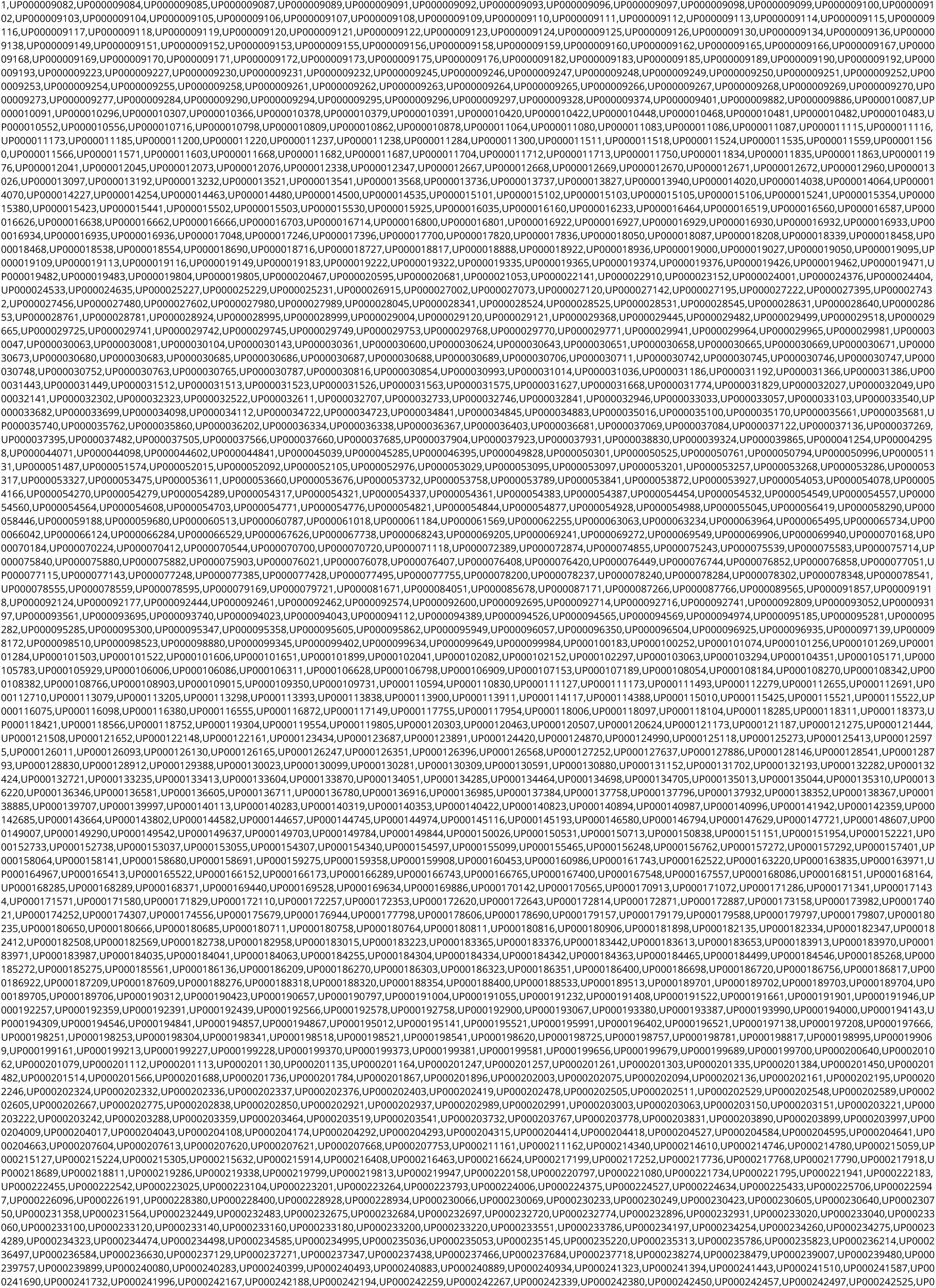
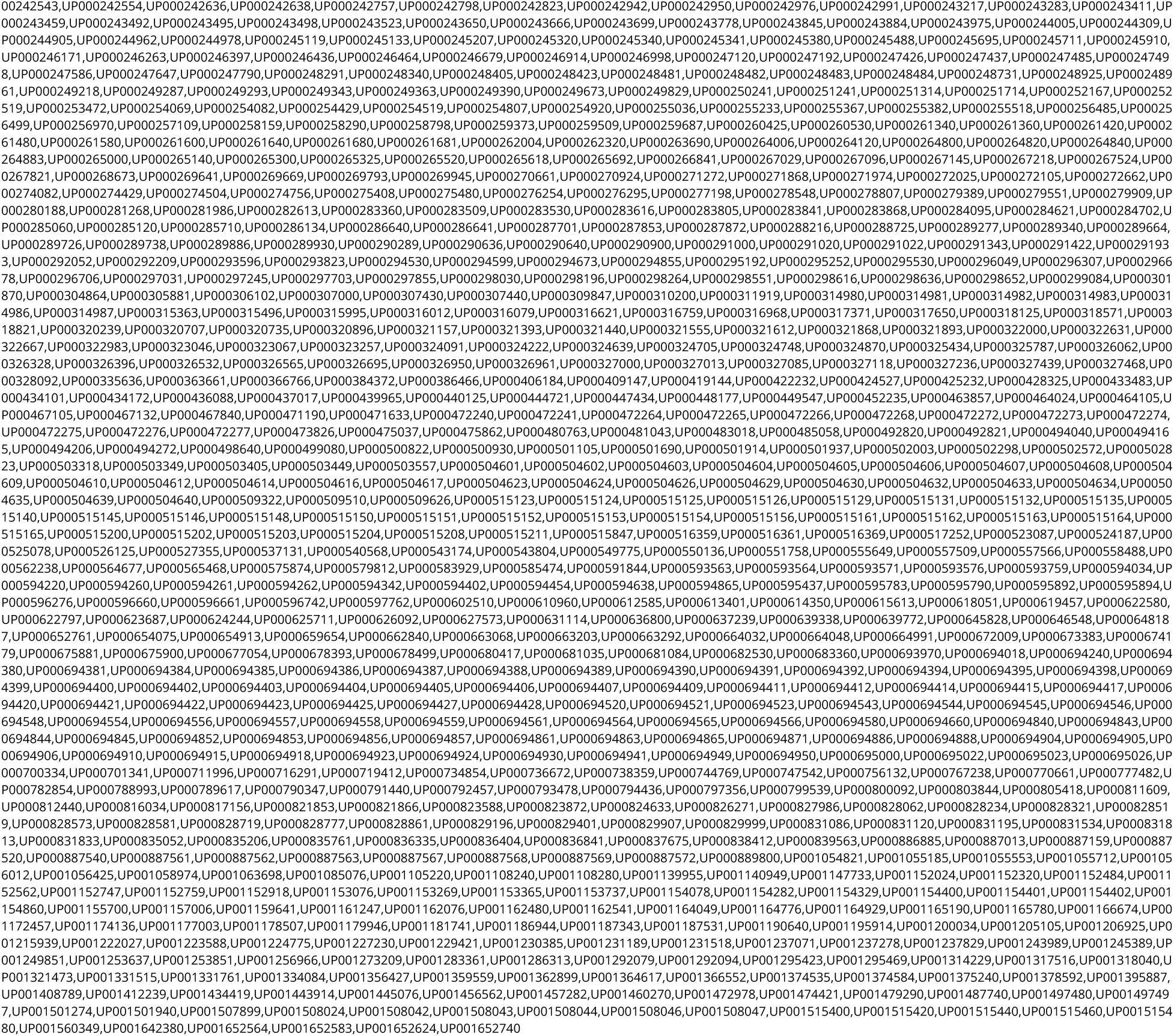

